# Systemic RNA interference-defective (SID) genes modulate dopaminergic neurodegeneration in *C. elegans*

**DOI:** 10.1101/2022.02.23.481573

**Authors:** Anthony L. Gaeta, J. Brucker Nourse, Karolina Willicott, Luke E. McKay, Candice M. Keogh, Kylie Peter, Shannon N. Russell, Shusei Hamamichi, Laura A. Berkowitz, Kim A. Caldwell, Guy A. Caldwell

## Abstract

The fine-tuning of gene expression is critical for all cellular processes; aberrations in this activity can lead to pathology, and conversely, resilience. As their role in coordinating organismal responses to both internal and external factors have increasingly come into focus, small non-coding RNAs have emerged as an essential component to disease etiology. Using Systemic RNA interference Defective (SID) mutants of the nematode *Caenorhabditis elegans*, deficient in endogenous gene silencing, we examined the potential consequences of dysfunctional epigenomic regulation in the context of Parkinson’s disease (PD). Specifically, the loss of either the *sid-1* or *sid-3* genes, which encode a dsRNA transporter and an endocytic regulatory non-receptor tyrosine kinase, respectively, conferred neuroprotection to dopaminergic neurons in an established transgenic *C. elegans* strain wherein overexpression of human α-synuclein (α-syn) from a chromosomally integrated multicopy transgene causes neurodegeneration. We further show that knockout of a specific microRNA, *mir-2,* attenuates α-syn neurotoxicity; suggesting that the native targets of *mir-2*-dependent gene silencing represent putative neuroprotective modulators. In support of this, we demonstrated that RNAi knockdown of multiple *mir-2* targets enhanced α-syn-induced dopaminergic neurodegeneration. Moreover, we demonstrate that *mir-2* overexpression originating in the intestine can induce neurodegeneration of dopaminergic neurons, an effect that was reversed by pharmacological inhibition of SID-3 activity. Interestingly, *sid-1* mutants retained *mir-2*-induced enhancement of neurodegeneration. Transcriptomic analysis of α-syn animals with and without a *sid-1* mutation revealed 27 differentially expressed genes with human orthologs related to a variety of diseases, including PD. Among these was *pgp-8*, encoding a P-glycoprotein-related ABC transporter. Notably, *sid-1*; *pgp-8* double mutants abolished the neurodegeneration resulting from intestinal *mir-2* overexpression. This research positions known regulators of small RNA-dependent gene silencing within a framework that facilitates mechanistic evaluation of epigenetic responses to exogenous and endogenous factors influencing dopaminergic neurodegeneration, revealing a path toward new targets for therapeutic intervention of PD.

**Author Summary:** The progressive death of neurons that produce the neurotransmitter dopamine is a clinical hallmark of Parkinson’s disease (PD). An integrated response to environmental and genetic factors leads to expression changes in specific genes and non-protein-coding regulatory molecules called dsRNAs that influence the pathology underlying PD. Here we report, for the first time in an animal model of PD, that mutations in genes encoding proteins which function in the cellular import of dsRNA protect dopamine neurons from degeneration. By generating a profile of individual genes affected when dsRNA transport is incapacitated, we established a foundation for the systematic analysis of their distinct contributions to neuroprotection. One subclass of dsRNAs, termed microRNAs, function analogously to a conductor of an orchestra by controlling hundreds of genes, simultaneously, to coordinate cellular processes. We identified a single microRNA that suppresses the activity of numerous genes in dopamine neurons and contributes to neurodegeneration. Genomic knockout of this deleterious microRNA, or prevention of its transport into dopamine neurons, unmasked a set of previously unreported neuroprotective proteins. This study supports a hypothesis whereby mechanisms that “fine tune” dopamine availability intersect with regulators of dsRNA transport to cooperatively maintain an optimal balance between neuronal activity, survival, and neurodegeneration.

## Introduction

The silencing of genes is a fundamental mechanism by which cells can dynamically regulate gene expression to meet the specific needs of an organism more precisely [1,2,3]. Being so important for organismal health, and potentially affecting all aspects of ecological fitness, it is no surprise that there are a multitude of ways that cells can execute this process. These include utilizing small, double-stranded RNAs (dsRNAs) such as microRNAs (miRNAs), piwi-interacting RNAs (piRNAs), and small interfering RNAs (siRNAs), to alter the expression of genes or change the epigenetic landscape of DNA that can indirectly lead to differential gene expression [4,5,6]. Gene silencing by dsRNAs can be accomplished by either full or partial binding to messenger RNA (mRNA) transcripts and subsequently suppressing or completely inhibiting the translation of the transcript into protein [2, 7]. Several organisms have evolved mechanisms to endogenously express and transport small dsRNAs in the coordinated regulation of gene expression, facilitating a rapid, dynamic, and adaptive means of organismal response to internal or external conditions [6, 8]. Researchers have also learned to exploit the exquisite specificity of this mechanism, using RNA interference (RNAi) to deplete target RNA transcripts and determine the functional implications of knocking down the activity of distinct genes. In this manner, a greater understanding of the respective impact that individual genes have as contributors to general organismal health and specific phenotypes, mechanisms, or pathways, has been substantially expedited through the application of RNAi [9,10,11,12,13]. Its revolutionary role in expanding our functional genomic toolkit aside, significant gaps remain in our basic understanding of RNAi and how it is effectuated through the organismal transmission of dsRNAs.

Small RNA-dependent changes in gene expression have been shown to contribute not only to beneficial processes such as stress responses, but also to pathology, where they have been revealed to be key factors in these processes [14,15,16,17,18]. *Caenorhabditis elegans*, possess a class of genes including *sid-1* and *sid-3*, designated Systemic RNA Interference Defective (SID), that encode proteins necessary for the organismal distribution and cellular transport of dsRNAs to trigger the selective targeting of transcripts for silencing [19, 20]. SID-1 is a transmembrane transporter protein that imports dsRNA into various cell types, allowing for RNA-mediated gene silencing [20,21,22]. Mutant *sid-1* animals are phenotypically characterized as being resistant to RNAi and lack the capacity to confer the systemic spread of dsRNA across the cells and tissues of this metazoan nematode [21]. SID-3 is a non-receptor tyrosine kinase that resides in the cytoplasm, exhibits binding activity to clathrin heavy chains, and prevents clathrin-dependent endocytosis, which ultimately prevents internalization of plasma membrane-localized proteins [23, 24]. The mammalian ortholog of SID-3, ACK1, functions as a form of “brake” on endocytosis, which when released leads to the endocytic recycling of plasma membrane proteins. In dopamine (DA) neurons, ACK1 has been shown to regulate the density of the transmembrane dopamine transporter (DAT) on neural membranes, thereby modulating DA reuptake from the synapse [25]. In an analogous scenario, *C. elegans* SID-3 is required for the efficient import of dsRNA into cells, as *sid-3* mutants exhibit a significant reduction in the capacity for environmental RNAi [23]. We posit that this is due to the involvement of SID-3 in maintaining baseline levels of SID-1 on cell surfaces, parallel to the activity of ACK1 in modulating DAT internalization.

In this study, transgenic *C. elegans* overexpressing a hallmark pathological gene product of Parkinson disease (PD), human α-synuclein (α-syn), are used to determine the effects of alterations in the systemic dsRNA uptake machinery on neuronal health and survival in the context of PD. Loss-of-function (*lof*) mutations in either *sid-1* or *sid-3*, were both found to protect DA neurons from the neurotoxic effects of transgenic multicopy α-syn overexpression in an established *C. elegans* model for the quantitative evaluation of DA neurodegeneration.

The collective outcomes of this research support a hypothesis that asserts factors influencing the systemic transport and cellular translocation of small dsRNAs represent previously undefined neuromodulators of DA neuron survival in the presence of proteotoxic stress resulting from α-syn overexpression in *C. elegans*. We examine this hypothesis through a combination of transcriptomic profiling of differential gene expression in *sid-1* mutants, as well through a combination of mutant analysis and selective overexpression of a specific *C. elegans* miRNA termed *mir-2* that modulates the progressive α-syn-induced DA neurodegeneration observed in PD. We report results of a systematic analysis of evolutionarily conserved and functionally validated targets of *mir-2* regulation for their contribution to a state of heightened neuroprotection that is observed in the absence of *mir-2* transcriptional repression. These experiments add support for prior associative genetic relationships with PD and reveal new targets of potential therapeutic significance. Additionally, we utilize tissue-specific transgene expression to evaluate miRNA-mediated, cell non-autonomous effects on the DA neurodegeneration induced by α-syn in defined genetic mutant backgrounds to establish relationships between specific transport proteins as functional gatekeepers of dsRNA-dependent gene silencing. Taken together, this research advances our mechanistic understanding of epigenetic factors influencing organismal dynamics that effectively alter the intrinsic threshold of neuroprotection to the proteotoxic dysregulation underlying PD.

## Results

### *sid-1* and *sid-3* mutants exhibit reduced dopaminergic neurodegeneration in a transgenic *C. elegans* α-syn model of PD

Evidence strongly suggests that SID-1 localization is regulated by SID-3 [23], which putatively acts to maintain SID-1 at plasma membranes and facilitates target gene silencing via dsRNA import by SID-1. To determine if endogenous gene silencing modulates dopaminergic neurodegeneration on its own, worms overexpressing GFP under the control of the DA neuron-specific *dat-1* promoter were crossed to *sid-1* mutants. The *sid-1(pk3321)* missense mutant has been shown to be defective in its normal function to “spread” or transmit dsRNA to other cells and tissues [21]. The *sid-1* mutation, and thus a loss of gene silencing ability, was found to have no significant effect on neurodegeneration at days 5, 7, and 10 post-hatching (Fig. 1A). To determine if endogenous gene silencing is involved in modulating α-syn-mediated DA neurodegeneration, transgenic nematodes overexpressing both GFP and human, wild-type α-syn exclusively in DA neurons were crossed to *sid-1* and *sid-3* mutants. The *sid-3 ok973* allele induces a deletion of 1,330 basepairs and an insertion of 12 basepairs, deleting exons 11 and 12, and part of exon 13 [26]. When GFP is solely expressed in the DA neurons, a negligible amount of neurodegeneration is observed (Fig. 1G), whereas co-expression with α-syn results in progressive degeneration of DA neurons that worsens with age (Fig. 1H). The *sid-1* mutation in the α-syn background did not significantly modulate neurodegeneration at day 5 or day 7 post-hatching, but significantly reduced neurodegeneration at day 10 post-hatching when compared to the α-syn background alone (Fig. 1B, Fig. 1I). This reveals an age-associated benefit conferred by *sid-1* mutants for DA neurons towards an increased capacity to resist the cellular stress posed by multicopy α-syn gene expression. In a reciprocal experiment, the neuroprotection bestowed by the *sid-1* mutation at day 10 post-hatching is lost when SID-1 activity is restored by selective *sid-1* overexpression in the DA neurons (Fig. 1C). Conversely, the *sid-3* mutation in the α-syn background significantly reduced neurodegeneration at all ages examined (days 5, 7, and 10 post-hatching) (Fig. 1D, Fig. 1J). When both *sid-1* and *sid-3* are mutant in the α-syn background, no significant change in neurodegeneration was observed at day 5 post-hatching, a time-point when the *sid-3* mutation independently exhibited significant neuroprotection, and when the *sid-1* mutation alone displayed no significant change in DA neurodegeneration (Fig. 1B). Moreover, this *sid-1*; *sid-3* double mutant exhibited robust neuroprotection at days 7 and 10 post-hatching (Fig. 1E, Fig. 1K), displaying a similar phenotype to the *sid-3* mutation alone at the same time points. Thus, the elimination or reduction of these established regulators of endogenous gene silencing leads to protection in an α-syn model of dopaminergic neurodegeneration (Fig. 1L).

**Fig. 1.**
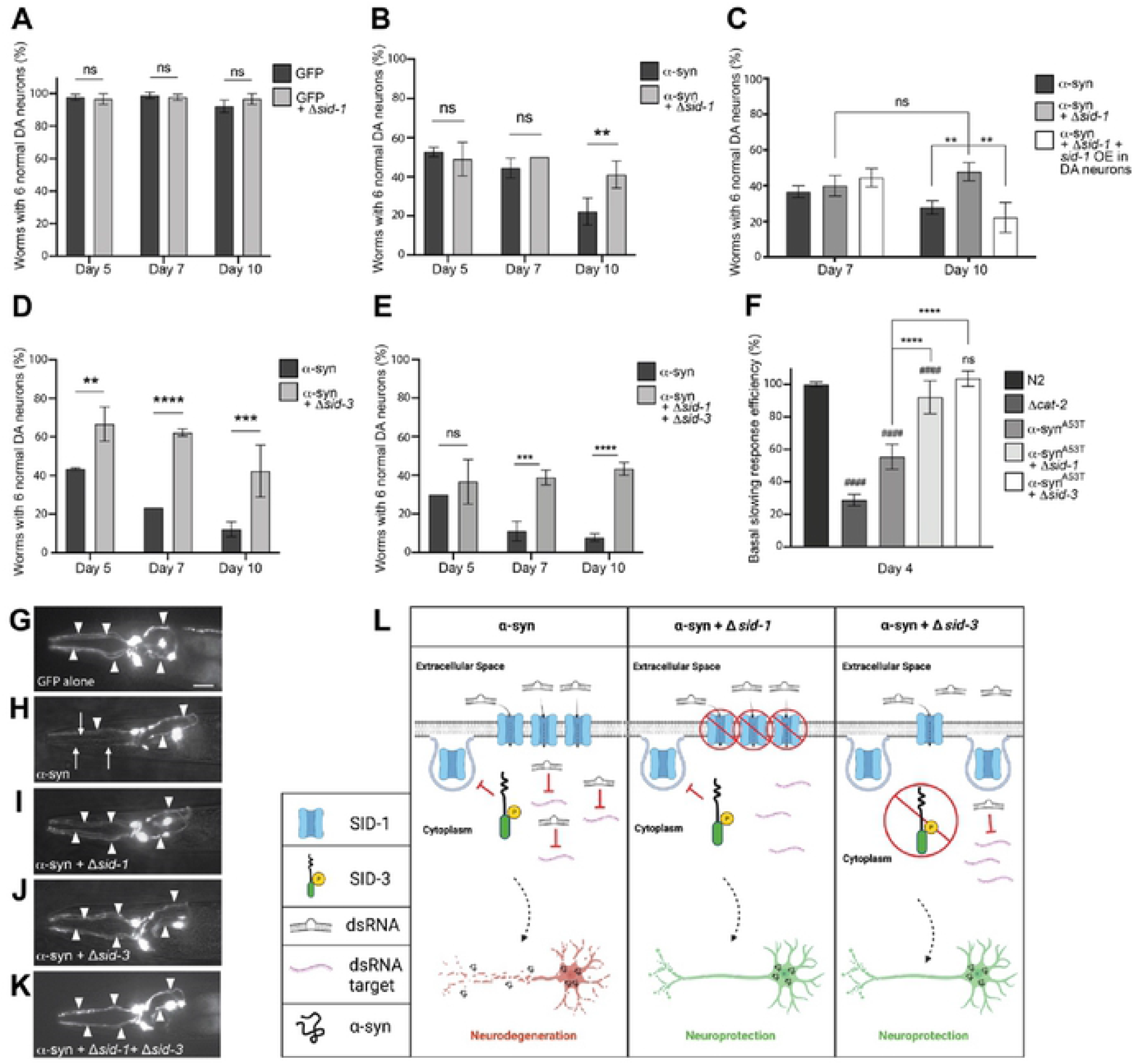
Loss of SID-1 and/or SID-3 functionality decreases dopaminergic neurodegeneration and improves DA neuron function. (**A**) Dopaminergic neurons were scored for degeneration on days 5, 7, and 10 post-hatching. Values represent mean ± S.D. (n= 30 worms per genotype per replicate, 3 independent replicates). Two-Way ANOVA with Šídák’s *post hoc* analysis was used to compare GFP alone (P*_dat-1_*::GFP, wild-type *sid-1*) to GFP + Δ*sid-1(pk3321)* at each time point; ns P ≥ 0.05. (**B**) Dopaminergic neurons were scored for degeneration on days 5, 7, and 10 post-hatching. Values represent mean ± S.D. (n= 30 worms per genotype per replicate, 3 independent replicates). Two-Way ANOVA with Šídák’s *post hoc* analysis was used to compare neurodegeneration in an α-syn model background (P*_dat-1_*::GFP + P*_dat-1_*::α-syn, wild-type *sid-1*), where α-syn is expressed in the dopaminergic neurons only, and in the α-syn worms with Δ*sid-1*, at each time point; ns P ≥ 0.05, ** P < 0.01. (**C**) Dopaminergic neurons scored for neurodegeneration on day 7 and 10 post-hatching. Values represent mean ± S.D. (n= 30 worms per genotype per replicate, 3 independent replicates). Two-Way ANOVA with Tukey’s *post hoc* analysis was used to compare all conditions to each other [an α-syn model when *sid-1* is wildtype, mutant, or mutant with *sid-1* overexpressed (OE) in DA neurons (P*_dat-1_*::*sid-1*)]; ns P ≥ 0.05, ** P < 0.01. (**D**) Dopaminergic neurons were scored for degeneration on days 5, 7, and 10 post-hatching. Values represent mean ± S.D. (n= 30 worms per genotype per replicate, 3 independent replicates). Two-Way ANOVA with Šídák’s *post hoc* analysis was used to compare α-syn alone to α-syn + Δ*sid-3(ok973)* at each time point; ** P < 0.01, *** P <0.001, **** P < 0.0001.(**E**) Dopaminergic neurons were scored for degeneration on days 5, 7, and 10 post-hatching. Values represent mean ± S.D. (n= 30 worms per genotype per replicate, 3 independent replicates). Two-Way ANOVA with Šídák’s *post hoc* analysis was used to compare α-syn alone to α-syn + Δ*sid-1;* Δ*sid-3* at each time point; ns P ≥ 0.05, *** P <0.001, **** P < 0.0001. (**F**) Basal Slowing Response Efficiency (BSR) assay, normalized to N2. *cat-2* mutants synthesize significantly less DA than N2 worms and serve as a positive control. Transgenic worms expressing a familial, mutant of α-syn, without GFP in the neurons, P*_dat-1_*::α-syn^A53T^, along *sid-1(pk3321)* and *sid-3(ok973)* mutant backgrounds, were used in this assay to quantify early changes in neurotransmission. Worms were tested on day 4 post-hatching. One-Way ANOVA with Tukey’s *post hoc* analysis was used to compare all conditions to each other. Symbols above bars are in comparison to N2 control; ns P ≥ 0.05, #### P < 0.0001. Symbols above brackets are in comparison to bars indicated. **** P < 0.0001. (**G-K**) These images represent the six anterior DA neurons in characteristic worms expressing GFP in the six head neurons on day 10 post-hatching. Arrowheads indicate intact DA neurons while arrows indicate degenerated DA neurons. Scale bar, 20 μm. (**G**) This animal expresses GFP only and displays a full complement of 6 anterior DA neurons. (**H**) GFP + α-syn expressed in DA neurons where the arrowheads indicate only 3 neurons are intact while 3 neurons are degenerating (arrows). (**I**) GFP + α-syn expressed in DA neurons in a *sid-1(pk3321)* mutant background; all 6 anterior DA neurons are intact. (**J**) GFP + α-syn expressed in DA neurons in a *sid-3(ok973)* mutant background; all 6 anterior DA neurons are intact. (**K**) GFP + α-syn expressed in a *sid-1(pk3321); sid-3(ok973)* double mutant background; all 6 anterior DA neurons are intact. (**L**) A tripartite illustration (created with Biorender.com) depicting an interpretation of the results obtained where the activity of SID-1 and/or SID-3 modulates dsRNA import, leading to differential effects on α-syn-induced DA neurodegeneration. The left pane displays a scenario where α-syn is expressed in DA neurons, while *sid-1* and *sid-3* are both wild-type: activated SID-3 functions as an endocytic brake, thereby preventing endocytosis of SID-1, which maintains the transport of dsRNAs that silence target genes in DA neurons, rendering them vulnerable to neurodegeneration. The middle pane displays the situation when α-syn is expressed in DA neurons of animals in which *sid-1* is mutant and *sid-3* is wild-type: here, the loss of SID-1 function results in reduced dsRNA transport, thereby leading to the neuroprotection from α-syn overexpression observed, as a logical consequence of attenuated gene silencing and the concomitant transcriptional upregulation of protective gene products. In the right pane, *sid-1* is wild-type and *sid-3* is mutant in the same neurotoxic α-syn background: when SID-3 is incapacitated by mutation the endocytosis of SID-1 is no longer blocked, resulting in diminished dsRNA import and a reduction in the silencing of neuronal target gene expression that leads to the enhanced neuroprotection observed experimentally.

### *sid-1* and *sid-3* mutants exhibit an improved neurobehavioral response in a transgenic *C. elegans* model of familial PD

Next, we wanted to determine if the reduced neurodegeneration observed in *sid-1* and *sid-3* mutant animals was also reflected by improved DA neurotransmission. To test this, Basal Slowing Response (BSR) assays were performed on *sid-1* and *sid-3* mutant animals. The BSR is a mechanosensory behavioral readout for DA neuron signaling and health and is dependent on the normal functionality of DA neurons [27]. Wildtype (N2) and *cat-2* mutant (DA deficient; tyrosine hydroxylase mutant) worms were used as controls to validate the assay, as N2 worms exhibit a normal BSR and *cat-2* mutant worms exhibit a defective BSR [27]. The *sid-1* and *sid-3* mutants were crossed into transgenic animals that overexpress a specific familial mutant form of α-syn (A53T) solely in the DA neurons. The A53T mutation increases the rate of α-syn oligomerization and leads to an early-onset, familial form of PD [28, 29]. Transgenic α-syn^A53T^ worms exhibited a defective BSR compared to N2 controls, as expected (Fig. 1F). Worms harboring either the *sid-1* or *sid-3* mutation in this α-syn background displayed a significantly improved BSR efficiency compared to the α-syn^A53T^ worms alone. The *sid-3* mutation rescued the BSR to such a degree that it was indistinguishable from wildtype (N2) animals (Fig. 1F). Taken together with their impact on neurodegeneration, these behavioral data support the notion that decreases in SID-1 and SID-3 activity robustly protect DA neurons from the neurotoxicity associated with excess α-syn, implying that loss and/or a reduction of endogenous RNA-mediated gene silencing is beneficial, in this context.

### Pharmacological inhibition of SID-3 diminishes the efficacy of exogenous RNAi and impact on DA neurodegeneration

Having demonstrated that α-syn-associated DA neurodegeneration displayed significant sensitivity to the presence or absence of SID-1, we set out to determine the consequences of attenuating the activity of SID-3 which, by extension of its known endocytic function, modulates levels of SID-1 in DA neurons and elsewhere in *C. elegans*. Although *sid-3* mutants have been shown to reduce the effectiveness of exogenous environmental RNAi in terms of systemic knockdown of corresponding genomic targets [23], it is unclear what effect reduction of SID-3 activity would have on neurodegeneration. The compound AIM-100 has been previously shown to inhibit the activity of ACK1 [30, 31], the human ortholog of SID-3. To determine if administration of AIM-100 could reduce the general effectiveness of environmental RNAi, GFP was knocked down via RNAi bacterial feeding. In these experiments, animals of a strain of *C. elegans*, HC46 [21], that overexpress nuclear-localized GFP in the large and readily observable body wall muscles, were reared on GFP-targeting RNAi bacteria, with or without AIM-100 (100 μM in 0.1% ethanol), and subsequently examined for GFP fluorescence. We predicted that AIM-100 would prevent GFP silencing via SID-3 inhibition. Indeed, animals that were fed control RNAi bacteria (empty vector) and not exposed to AIM-100 (ethanol solvent only) displayed robust expression of GFP in the body wall muscles (Fig. 2A). When GFP was knocked down in animals not exposed to AIM-100, the GFP fluorescence was silenced in the body wall muscles to a large degree, as expected (Fig. 2B). When GFP was targeted for knockdown in worms that were exposed to AIM-100, GFP was unable to be silenced in the body wall muscles to the same extent and exhibited noticeably brighter and more abundant fluorescence (Fig. 2C). Therefore, a drug-induced reduction in SID-3 activity via AIM-100 diminishes the efficacy of exogenous RNAi knockdown, consistent with prior studies targeting GFP in the body wall muscles in genetic mutants of *sid-3* [23].

**Fig. 2.**
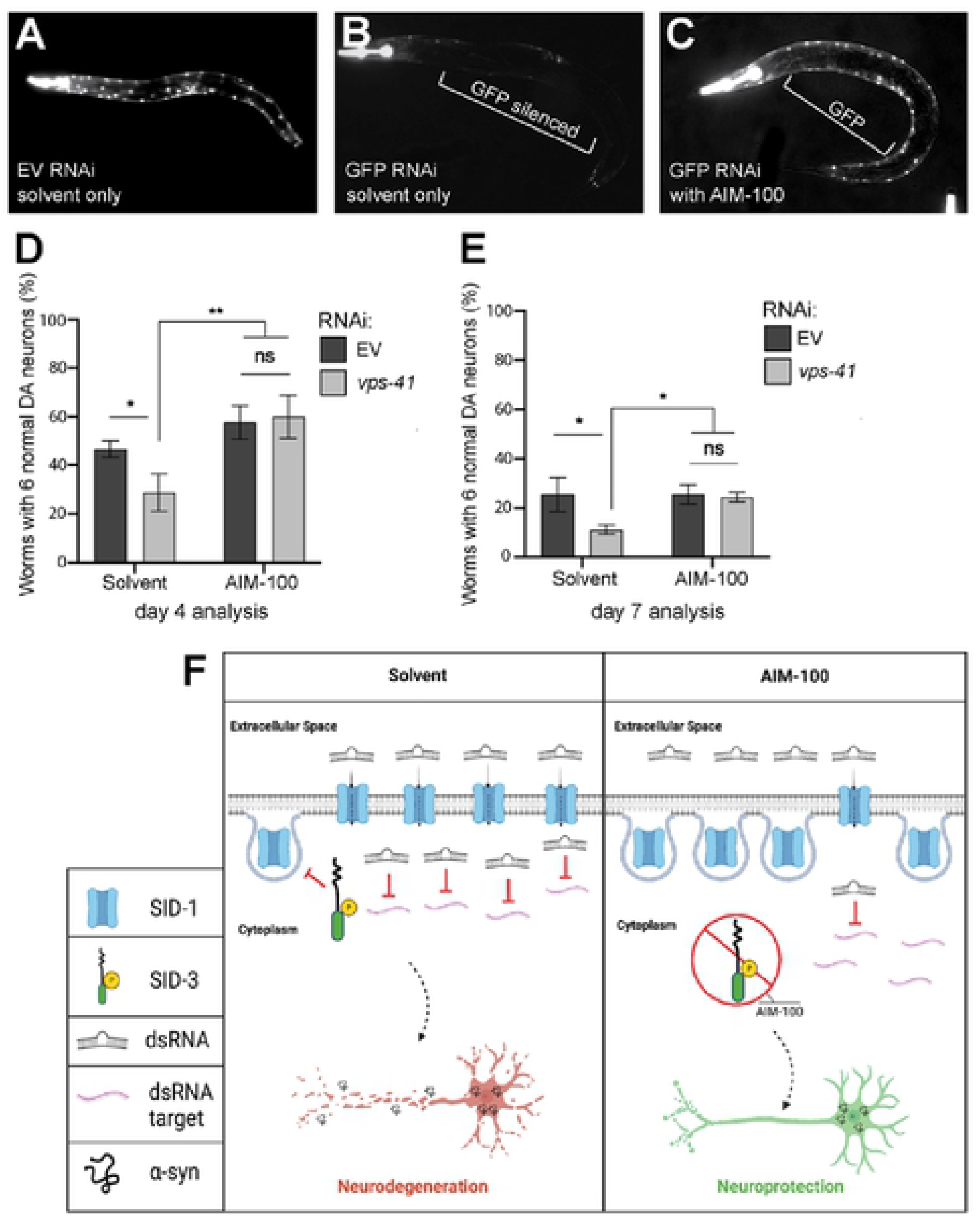
AIM-100 reduces dsRNA silencing in body wall muscle and decreases dopaminergic neurodegeneration. (**A-C**) RNAi in worms overexpressing GFP in body wall muscle cells, with and without the SID-3 inhibitor AIM-100 (100 μM) in 0.1% ethanol solvent. Worms were tested on day 5 post-hatching. (**A**) Empty vector (EV) RNAi in worms overexpressing GFP in body wall muscle cells exposed. (**B**) GFP RNAi in worms overexpressing GFP in body wall muscle cells. (**C**) GFP RNAi in worms overexpressing GFP in body wall muscle cells and administered AIM-100. (**D, E**) Dopaminergic neurodegeneration levels in an RNAi-sensitive α-syn model background designed to allow knockdown of genes solely in DA neurons (strain UA196) and analyzed at days 4 and 7 post-hatching. Worms were either exposed to the 0.1% ethanol solvent control or AIM-100 (100 μM), and within each of these conditions, mock RNAi (empty vector) or *vps-41* (positive control) RNAi was performed. Values represent mean ± S.D. (n= 30 worms per genotype per replicate, 3 independent replicates). Two-way ANOVA with Tukey’s *post hoc* analysis was used to compare all conditions to each other; ns P ≥ 0.05, * P < 0.05, ** P < 0.01. Values represent mean ± S.D. (n= 30 worms per genotype per replicate, 3 independent replicates). (**F**) An illustration (created with Biorender.com) depicting model scenarios for the import of dsRNA (targeting *vps-41,* an established control for DA neurodegeneration when knocked down) into DA neurons via SID-1 in transgenic animals expressing α-syn, either in the presence (left pane) or absence (right pane) of treatment with the selective inhibitor of SID-3 activity, AIM-100. The left pane represents SID-3 actively blocking the endocytosis of SID-1 in the absence of AIM-100 (solvent only), therefore maintaining dsRNA transport for the silencing of target genes, such as *vps-41*, which enhances DA neurodegeneration when knocked down. In contrast, the right pane displays a scenario that explains the observed inhibition of SID-3 by AIM-100 treatment seen when α-syn is expressed in DA neurons; in this scenario AIM-100 reduces the activity of SID-3 and allows for endocytosis of SID-1, thereby resulting in less SID-1 on cell surfaces and reduced dsRNA import into cells. This leads to the stability of endogenous *vps-41* transcripts and enhanced resistance to a-syn-induced neurodegeneration observed, in accordance with the established function of VPS-41.

To determine if AIM-100 administration could reduce the effectiveness of exogenous RNAi in DA neurons overexpressing α-syn, we used an α-syn transgenic strain that enables the selective knockdown of genes only in the DA neurons. This strain (UA196) overexpresses human wild-type α-syn from a chromosomally integrated multicopy transgene in a *sid-1* mutant background, and concomitantly overexpresses *sid-1* from the *dat-1* promoter thereby restoring delimited RNAi sensitivity in just the DA neurons [32]. These worms were fed either empty vector (negative control) RNAi bacteria or RNAi bacteria producing dsRNA targeting *vps-41*, an internal control which faithfully enhances neurodegeneration due to a reduction in endolysosomal trafficking, an essential process for DA neuron health [33, 34]. We carried out these RNAi assays with the addition of AIM-100, predicting that worms grown on *vps-41* RNAi bacteria in the presence of AIM-100 would display significantly less neurodegeneration compared to RNAi in the solvent-treated group. At day 4 post-hatching, knockdown of *vps-41* significantly enhanced neurodegeneration compared to empty vector RNAi controls, in the solvent control group (Fig. 2D). However, knockdown of *vps-41* did not enhance neurodegeneration in the AIM-100-treated group (Fig. 2D). Very similar results were obtained when the same experiment was performed on day 7 post-hatching (Fig. 2E). In all, these results indicate that inhibition of SID-3 function by AIM-100 reduces the effectiveness of exogenous RNAi in both a non-neuronal tissue (body wall muscles) and in DA neurons (Fig. 2F). Given that AIM-100 is established as both a potent and selective inhibitor of mammalian ACK1 [35] these data on the worm ortholog, SID-3, indicate that this non-receptor tyrosine kinase represents a putative druggable target to modulate neurodegeneration through endocytic regulation of small RNA uptake.

### Modulation of α-syn-induced DA neurodegeneration by *mir-2*

Our previous findings revealed that a decrease in SID-3 function hindered RNAi when dsRNA was produced from exogenous sources (bacterial feeding). Next, we wanted to determine if attenuating SID-3 activity could similarly impede gene silencing resulting from a naturally occurring miRNA in *C. elegans*. Based on pre-existing database information on experimentally validated targets and the availability of a knockout mutation, we selected the microRNA, *mir-2*, for evaluation. When endogenous *mir-2* is mutant, protection is observed in the α-syn-induced model of DA neurodegeneration at both days 5 and 7 post-hatching (Fig. 3A). This suggests that the absence of epigenetic silencing of target gene expression by *mir-2* is beneficial for DA neurons in overcoming the stress from α-syn. It therefore follows that, in transgenic nematodes engineered to overexpress the *mir-*2 pre-miRNA exclusively in the DA neurons, along with α-syn, an increase in neurodegeneration was observed when compared to the α-syn background alone (Fig. 3B).

**Fig. 3.**
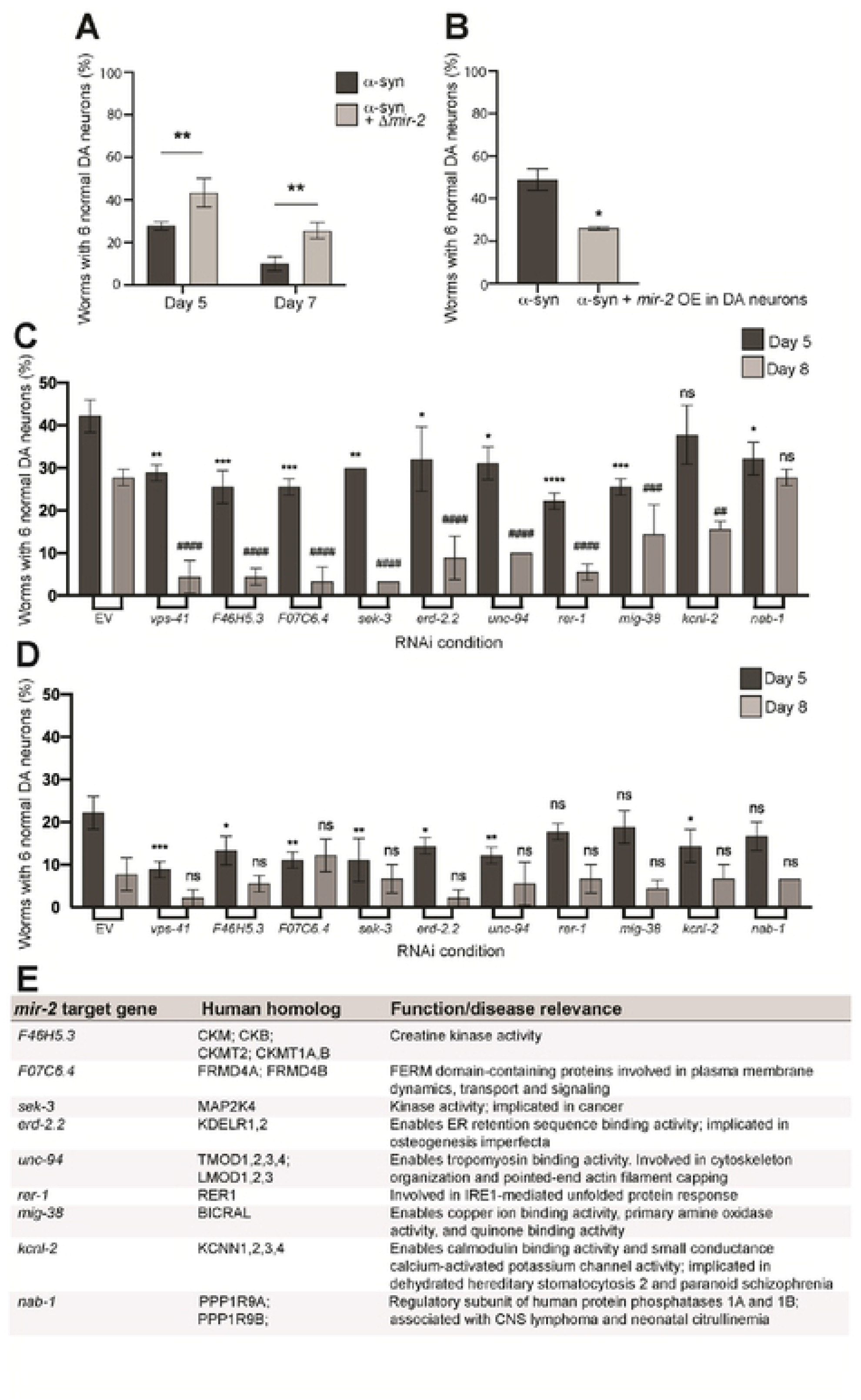
Modulating *mir-2*-dependent suppression of target gene expression levels alters α-syn-induced dopaminergic neurodegeneration. (**A**) Dopaminergic neurons were scored for degeneration on days 5 and 7 post-hatching. Values represent mean ± S.D. (n= 30 worms per genotype per replicate, 3 independent replicates). Two-Way ANOVA with Šídák’s *post hoc* analysis was used to compare α-syn alone α-syn + Δ*mir-2(gk259)* at each time point; ** P < 0.01. (**B**) Dopaminergic neurons were scored for degeneration on day 4 post-hatching. Values represent mean ± S.D. (n= 30 worms per genotype per replicate, 3 independent replicates). Unpaired, two-tailed Student’s *t*-test was used to compare α-syn with *mir-2* overexpression (OE) in dopaminergic neurons (P*_dat-1_*::*mir-2*) to α-syn only controls; * P < 0.05. (**C, D**) Dopaminergic neurons were scored for degeneration on days 5 and 8 post-hatching. RNAi was performed in a RNAi sensitive α-syn model with a *mir-2(gk259)* mutant background (**C**) and in the RNAi sensitive α-syn model that is *mir-2* wildtype (D). Nine validated targets of miR-2 that have human orthologs were knocked down, along with a positive control, *vps-41*. Values represent mean ± S.D. (n= 30 worms per genotype per replicate, 3 independent replicates). replicate, 3 independent replicates). One-Way ANOVA with Dunnett’s *post hoc* analysis was used to compare RNAi knockdowns to empty vector (EV) controls at day 5 and 8 post-hatching. For day 5 analyses, asterisks (*) above bars indicate comparisons of each individual knockdown to EV; ns P ≥ 0.05, * P < 0.05, ** P < 0.01, *** P <0.001, **** P < 0.0001. For day 8 analyses, pound signs (#) above bars indicate comparisons to EV; ns P ≥ 0.05, ## P < 0.01, ### P <0.001, #### P < 0.0001. (**E**) A list of the nine targets knocked down via RNAi in (**C**) and (**D**), their human homologs, and function.

### RNAi knockdown of conserved targets of *mir-*2 enhances α-syn-induced DA neurodegeneration

To gain clarity as to whether the observed effects on neurodegeneration observed from *mir-2* modulation could be attributed to gene silencing from within the DA neurons, individual target genes *of mir-2* were knocked down via RNAi. Only targets of *mir-2* that have human orthologs <u>and</u> have been previously validated as “bona fide” via a documented expression change (by RNAseq and/or qPCR in Wormbase) were selected for knockdown. These 9 genes code for proteins involved in a range of biological processes, including the unfolded protein response, cytoskeleton reorganization, and metal ion binding activity (Fig. 3E). We reasoned that RNAi knockdown of these targets specifically in the DA neurons <u>and</u> in a *mir-2* mutant background, where their baseline of expression would not be repressed, would uncover individual contributors to the neuroprotection we observed in the absence of *mir-2* (Fig. 3C). For this analysis, we employed the α-syn strain that allows for DA-neuron specific RNAi (UA196; used in Fig. 2D and E) after crossing the strain to the *mir-2* knockout mutant. The outcomes of this effort indicated that the knockdown of 8 of 9 validated *mir-2* targets resulted in an increased percentage of animals exhibiting DA neurodegeneration at day 5 post-hatching; a greater severity of degeneration was observed in these same targets by day 8 post-hatching (Fig. 3C). These targets were also knocked down in UA196, which was wild-type for *mir-2* (Fig. 3D). In this background, the knockdown of 6 of 9 validated *mir-2* targets resulted in an enhancement of neurodegeneration at day 5 post-hatching, whereas none of the knockdown of the *mir-2* targets had any significant effect on neurodegeneration at day 8 post-hatching, likely due to the extent of neurodegeneration observed at this time point in the controls (Fig. 3D). These results are consistent with the interpretation that endogenous suppression of these genes by *mir-2* restricts their inherent neuroprotective capacity. Although these data reveal the cell autonomous effects *of mir-2* in DA neurons, we wanted to further explore the prospect that effectors of miRNA transport might also modulate α-syn-induced neurodegeneration through cell non-autonomous overexpression of this specific miRNA from the worm intestine.

### DA neurodegeneration by α-syn is enhanced by cell non-autonomous overexpression of *mir-2* and is modulated by AIM-100

Having determined that overexpression of *mir-2* in the DA neurons directly affected their survival in the presence of α-syn, we further explored the impact of *mir-2* produced from an indirect cellular source. We generated a strain which overexpressed *mir-2* (pre-miRNA) in the intestinal compartment (gut) under the control of the intestinal-specific promoter, *ges-1*, and crossed it into *C. elegans* expressing human, wild-type α-syn in the DA neurons (Fig. 4A). Unexpectedly, the tissue-specific overexpression of *mir-2* delimited to the worm gut led to a significant enhancement of DA neurodegeneration when compared to the α-syn background alone (Fig. 4A, Fig. 4H). To ascertain whether intestinal *mir-2* overexpression was sufficient to induce this enhancement of α-syn-induced neurodegeneration and that it was not dependent on the presence of an increase of endogenous *mir-2*, this strain was crossed into the *mir-2* knockout mutant previously examined (Fig. 3A). Interestingly, these animals also exhibited an enhancement of neurodegeneration when compared to the α-syn background alone, akin to wild-type endogenous *mir-2* (Fig. 4A). It is also important to note that the enhancement of neurodegeneration due to overexpression of *mir-2* appears to overshadow the inherent neuroprotective effect of the *mir-2* mutation itself. These results showcase the potency of *in vivo* epigenetic regulation by microRNAs across cellular boundaries in an intact metazoan. Specifically, the overexpression of *mir-2* from within the gut of *C. elegans* exerts a robust, cell non-autonomous effect on the survival of DA neurons already sensitized to neurodegeneration by α-syn-induced neurotoxicity. In a theoretical variation on this scenario, an analogous effect might result from a microbiome-based source of dsRNA or inducer of host changes in small RNA expression.

**Fig. 4.**
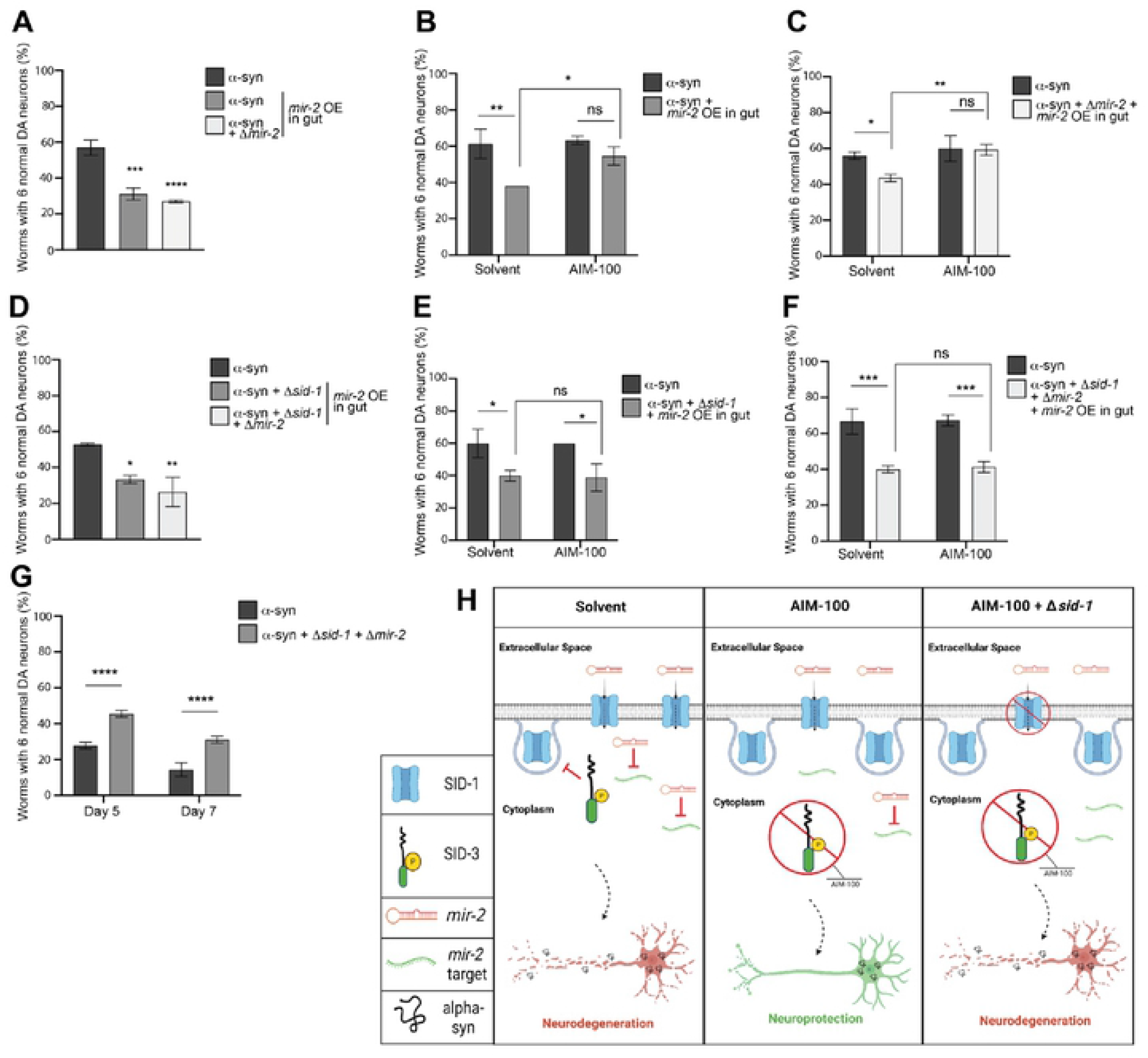
Intestinal overexpression of *mir-2* enhances dopaminergic neurodegeneration and is attenuated by AIM-100 when *sid-1* is wild-type. (**A, D**) Dopaminergic neurons were scored for neurodegeneration at day 4 post-hatching in an α-syn model background with *mir-2* OE in the intestine (P*_ges-1_*::*mir-2*). In (A), *mir-2(gk259)* is examined (mutant and wildtype) while in (**D**), both *sid-1(pk3321)* and *mir-2(gk259)* mutants and wildtype are examined. Values represent mean ± S.D. (n= 30 worms per genotype per replicate, 3 independent replicates). One-Way ANOVA with Dunnett’s *post hoc* analysis was used to compare worm classes where *mir-2* is OE in the intestine (P*_ges-1_*::*mir-2*) to the α-syn (with or without *sid-1* mutation) worm classes; * P < 0.05, ** P < 0.01, *** P <0.001, **** P < 0.0001. (**B-F**) Dopaminergic neurons were scored for degeneration on day 4 post-hatching. Values represent mean ± S.D. (n= 30 worms per genotype per replicate, 3 independent replicates). Worms were either exposed to the 0.1% ethanol solvent control or AIM-100 (100 μM), and the α-syn control model or the α-syn model with *mir-2* OE in the intestine (P*_ges-1_*::*mir-2*) alone (**B**), with *mir-2(gk259)* (**C**), *sid-1(pk3321)* (**E**), or both *sid-1(pk3321)* and *mir-2(gk259)* mutants (**F**) were examined. Two-way ANOVA with Tukey’s *post hoc* analysis was used to compare all conditions to each other; ns P ≥ 0.05, * P < 0.05, ** P < 0.01, *** P <0.001. (**G**) Dopaminergic neurons were scored for degeneration on days 5 and 7 post-hatching. Values represent mean ± S.D. (N = 3; n = 30 worms, per genotype). Two-Way ANOVA with Šídák’s *post hoc* analysis was used to compare α-syn expressed in neurons to α-syn + *sid-1(pk3321)* + *mir-2(gk259)* at each time point; **** P < 0.0001. (H) An illustrative triptych (created with Biorender.com) representing an interpretation of the results obtained for SID-3 regulation of *mir-2* import, modeling the differential impact of α-syn-induced DA neurodegeneration. The triptych displays conditions in which α-syn is expressed in DA neurons and *mir-2* is overexpressed (OE) in the worm intestine. In the left panel, *sid-1* and *sid-3* are wild-type, and SID-3 is depicted as inhibiting the endocytosis of SID-1, thus maintaining wild-type SID-1 levels for transport of *mir-2* and suppress the expression of its target genes, leaving DA neurons vulnerable to α-syn-induced neurodegeneration. The middle pane depicts these same animals treated with the potent and selective SID-3 inhibitor, AIM-100. When SID-3 is inhibited by AIM-100, evidence suggests that endocytosis of SID-1 is increased, resulting in a decrease of *mir-2* import into cells, and conversely increasing the abundance of upregulated transcripts that contribute to the DA neuroprotection observed. The right pane reflects the same scenario as does the middle, except under conditions where *sid-3* is wild-type*, sid-1* is mutant, and SID-3 is inhibited by AIM-100. If SID-3 activity is inhibited, increased internalization SID-1 from cell surfaces would be predicted, along with a corresponding decrease of *mir-2* import would be expected, aside from the fact that any SID-1 on cell surfaces is non-functional due to mutation. Strikingly, although *mir-2* import into cells via SID-1 is eliminated in this latter example, an enhancement of α-syn-induced neurodegeneration is still observed experimentally. This result suggests that an alternate mechanism of dsRNA uptake may exist and is revealed in the absence of functional SID-1.

To determine if the previously observed neuroprotective effects of SID-3 inhibition could influence the enhanced neurodegeneration induced by intestinal *mir-2* overexpression, we administered AIM-100 to these animals. In the solvent control group, *mir-2* overexpression in the gut enhanced neurodegeneration in accordance with previous results (Fig. 4B). In contrast, animals in the AIM-100-treated group no longer exhibited enhanced neurodegeneration as an outcome of *mir-2* overexpression (Fig. 4B). A very similar outcome was observed when these same strains were crossed into the *mir-2* mutant background: *mir-2* overexpression was no longer able to enhance neurodegeneration in the AIM-100 group (Fig. 4C). The observation that intestinal *mir-2* overexpression increases α-syn-mediated neurodegeneration but is unable to do so when AIM-100 is administered, suggests that the effects of intestinal *mir-2* overexpression are at least partly dependent on SID-3 function in blocking endocytic recycling of plasma membrane proteins, such as SID-1. Likewise, the neuroprotection observed in response to SID-3 inhibition by AIM-100 might be attributable to a decrease in *mir-2* transport, thereby limiting silencing of its targets (Fig. 4H).

To directly evaluate the effect of α-syn-induced DA neurodegeneration within the context of *sid-1* loss in worms overexpressing *mir-2* within the intestine, these animals were crossed into the *sid-1* mutant background, and DA neurons were scored for neurodegeneration (Fig. 4D). Surprisingly, when either *sid-1* mutants or *sid-1; mir-2* double mutants were crossed into *C. elegans* overexpressing *mir-2* from within the intestine (α-syn*^mir-2^* ^OE gut^), the increased neurodegenerative effect previously observed remained significant (Fig. 4D). This is especially noteworthy since *sid-1; mir-2* double mutants in the α-syn background are robustly neuroprotective compared to α-syn only controls, indicating that the neuroprotection conferred by these combined mutations is insufficient to combat neurodegeneration resulting from overexpression of *mir-2* in the intestine (Fig. 4G).

To determine if pharmacological inhibition of SID-3 impacted this observed cell non-autonomous effect on DA neurodegeneration in a manner independent of SID-1, we exposed either *sid-1* mutants or *sid-1; mir-2* double mutants in the α-syn*^mir-2^* ^OE gut^ background to AIM-100. Both these sets of *sid-1*-deficient animals exhibited significantly enhanced neurodegeneration when compared to α-syn worms alone in the solvent controls (Fig. 4E, Fig. 4F). When these same animals were exposed to AIM-100, intestinal overexpression of *mir-2* still led to significant enhancement of α-syn-induced DA neurodegeneration (Fig. 4E, Fig. 4F). These results collectively suggest that the DA neurodegeneration observed reflects the import of *mir-2* into DA neurons and is a potential consequence of the subsequent silencing of *mir-2* target genes. Significantly, the increased neurodegeneration observed in response to *mir-2* overexpression in the intestine occurs in the absence of SID-1 transporter function and SID-3 inhibition by AIM-100 (Fig. 4H).

### Transcriptomic comparison of transgenic α-syn animals with and without *sid-1*

We have shown that *sid-1* and *sid-3* mutants promote the ability of DA neurons to combat α-syn-associated neurotoxicity. To determine which genes, pathways, or biological processes may be contributing to the favorable effect on DA neurons observed in *sid-1* mutants, a comparative transcriptomic analysis was performed on our transgenic α-syn (wild-type, multicopy, human) animals with and without the *sid-1* mutation. This analysis revealed a total of 84 Differentially Expressed Genes (DEGs), 74 of which were upregulated in α-syn worms with the *sid-1* mutation and 10 of which were downregulated, when compared to the α-syn background strain containing functional *sid-1* (Fig. 5A). This bias towards DEGs being upregulated in animals with the *sid-1* mutation could be expected, since the expression of target genes normally suppressed by endogenous dsRNAs are no longer silenced in the absence of the SID-1 transporter.

**Fig. 5.**
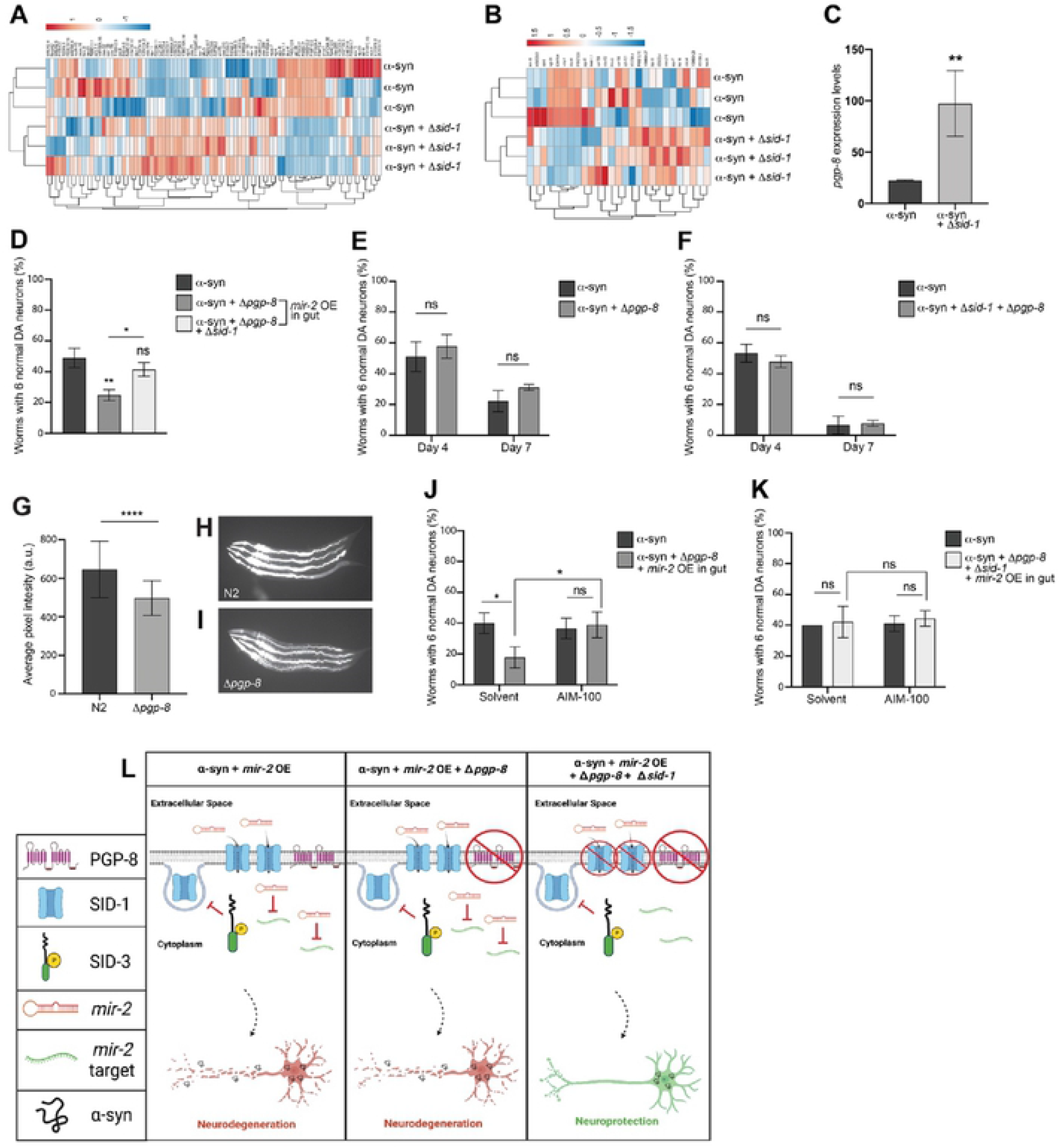
Loss of both SID-1 and PGP-8 functionality reduces *mir-2* OE-induced enhancement of neurodegeneration. (**A, B**) Heatmaps depicting regulatory status of the Differentially Expressed Genes (DEGs) (columns) from the transcriptomic analysis comparing α-syn and α-syn + Δ*sid-*1 groups (rows). Both the α-syn and α-syn + Δ*sid-1* groups constitute 3 replicates each. RNA was isolated at day 6 post-hatching. (**A**) Heatmap of all 84 DEGs associated with this analysis while be (**B**) depicts the regulatory status of the 27 DEGs with human orthologs. (**C**) *pgp-8* mRNA transcript levels in the α-syn background alone and the α-syn background with the *sid-1* mutation, identified from the transcriptomic analysis between the α-syn and α-syn + Δ*sid-1* groups. Both groups constitute 3 replicates each. The DESeq2 program with the Benjamini-Hochberg method to obtain adjusted p-values (Novogene Corporation, Inc.); ** P(adjusted) < 0.01. (**D**) Dopaminergic neurons were scored for neurodegeneration at day 4 post-hatching in an α-syn model background with *mir-2* OE in the intestine (P*_ges-1_*::*mir-2*), with a Δ*pgp-8(ok2489)* mutant and Δ*pgp-8(ok2489);* Δ*sid-1(pk3321)* double mutant. Values represent mean ± S.D. (n= 30 worms per genotype per replicate, 3 independent replicates). One-Way ANOVA with Tukey’s *post hoc* analysis was used to compare all conditions to each other; Symbols above bars are in comparison to α-syn only controls while the straight line is a comparison of the bars indicated. ns P ≥ 0.05, * P < 0.05, ** P < 0.01. (**E**, **F**) Worms expressing the α-syn control with *pgp-8(ok2489)* (**E**) or *pgp-8(ok2489); sid-1(pk3321)* (**F**) were analyzed for dopaminergic neurodegeneration at days 4 and 7 post-hatching. Values represent mean ± S.D. (n= 30 worms per genotype per replicate, 3 independent replicates). Error bars indicate standard deviation. ns P ≥ 0.05, by Two-Way ANOVA with Šídák’s *post hoc* analysis to compare α-syn alone with the mutant displayed in the same graph. (**G** - **I**) N2 wild-type and *pgp-8(ok2489)* mutant worms following a 48-hour exposure to 10 μM of rhodamine 123. (**G**) Average pixel intensity was measured 48 hours post-hatching. Values represent mean ± S.D. (n= 30 worms per genotype per replicate, 3 independent replicates). Unpaired, two-tailed Student’s *t*-test was used to compare N2 to *pgp-8(ok2489)*; **** P < 0.0001. Representative images of N2 (**H**) and *pgp-8* mutant (**I**) worms exhibiting fluorescence due to rhodamine 123 treatment. (**J, K**) Dopaminergic neurons were scored for neurodegeneration at day 4 post-hatching. Values represent mean ± S.D. (n= 30 worms per genotype per replicate, 3 independent replicates). Worms expressing the α-syn control or the α-syn with *mir-2* OE in the intestine (P*_ges-1_*::*mir-2*) with *pgp-8(ok2489)* (**J**) or *pgp-8(ok2489); sid-1(pk3321)* (**K**) were either exposed to the 0.1% ethanol solvent control or AIM-100 (100 μM). Two-way ANOVA with Tukey’s *post hoc* analysis was used to compare all conditions to each other; ns P ≥ 0.05, * P < 0.05. (**L**) An illustrative triptych illustration (created with Biorender.com) reflecting an interpretation of the experimentally observed impact on α-syn-induced dopaminergic neurodegeneration caused by intestinal overexpression of *mir-*2 in either a *pgp-8* knockout mutant or a *pgp-8*; *sid-1* double mutant. In the left pane, SID-3 blocks the endocytosis of SID-1 on cell membranes, maintaining ample SID-1 on cell surfaces to transport *mir-2* into cells, allowing for *mir-2* target gene silencing and leading to enhanced neurodegeneration. The middle pane displays the situation when α-syn is expressed in DA neurons and *mir-2* is OE in the intestine, while *sid-1* is wild-type and *pgp-8* is mutant. In this model, SID-3 blocks the endocytosis of SID-1 on the plasma membrane, resulting in the transport of *mir-2* for target gene silencing and the enhanced neurodegeneration observed experimentally. It is presumed that a loss of PGP-8 function does not influence *mir-2* import into cells, since SID-1 remains functional. The right pane is a model for when both *sid-1* and *pgp-8* are mutant in animals expressing α-syn in the DA neurons, with *mir-2* being overexpressed in the intestine. Here we hypothesize that *mir-2* might not be imported into cells either due to a mutation that renders SID-1 non-functional, or by the enigmatic process by which PGP-8 may influence *mir-2* import into cells, also associated with a mutation rendering it non-functional. In either case, less or no *mir-2* import into cells results, and instead, leads to an infusion of target gene transcription that contributes to the observed neuroprotection.

Out of the 84 DEGs, 27 had human orthologs, 22 of which were upregulated in *sid-1* mutants, and 5 of which were downregulated in *sid-1* mutants (Fig. 5B). These DEGs encode products predicted to function in a myriad of biological processes, including transmembrane transport, metabolism, cell signaling, stress response pathways, transcriptional regulation, proteolysis, and neuronal function (Table 1). From the 27 DEGs with human homologs, 18 (67%) were found to have a prior association to human diseases (Table 1). Notably, 9 of these DEGs have a previously identified association with PD in some way (33%); these include *clec-4*, *clec-52*, *dop-4*, *ugt-36*, *hmit-1.1*, *F49C12.6*, *F49E12.10*, *W07B8.4*, and *H39E23.3*. Interestingly, 4 other DEGs (*col-159*, *fmo-3, ech-1.1,* and *E04F6.4*) have a prior relationship to Alzheimer’s disease (AD) (Table 1). This candidate enrichment provides increased confidence that our strategy successfully informs us about mechanistic processes impacted by *sid-1*, and reveals specific, evolutionarily conserved gene products that contribute to neuroprotection.

**Table 1.**
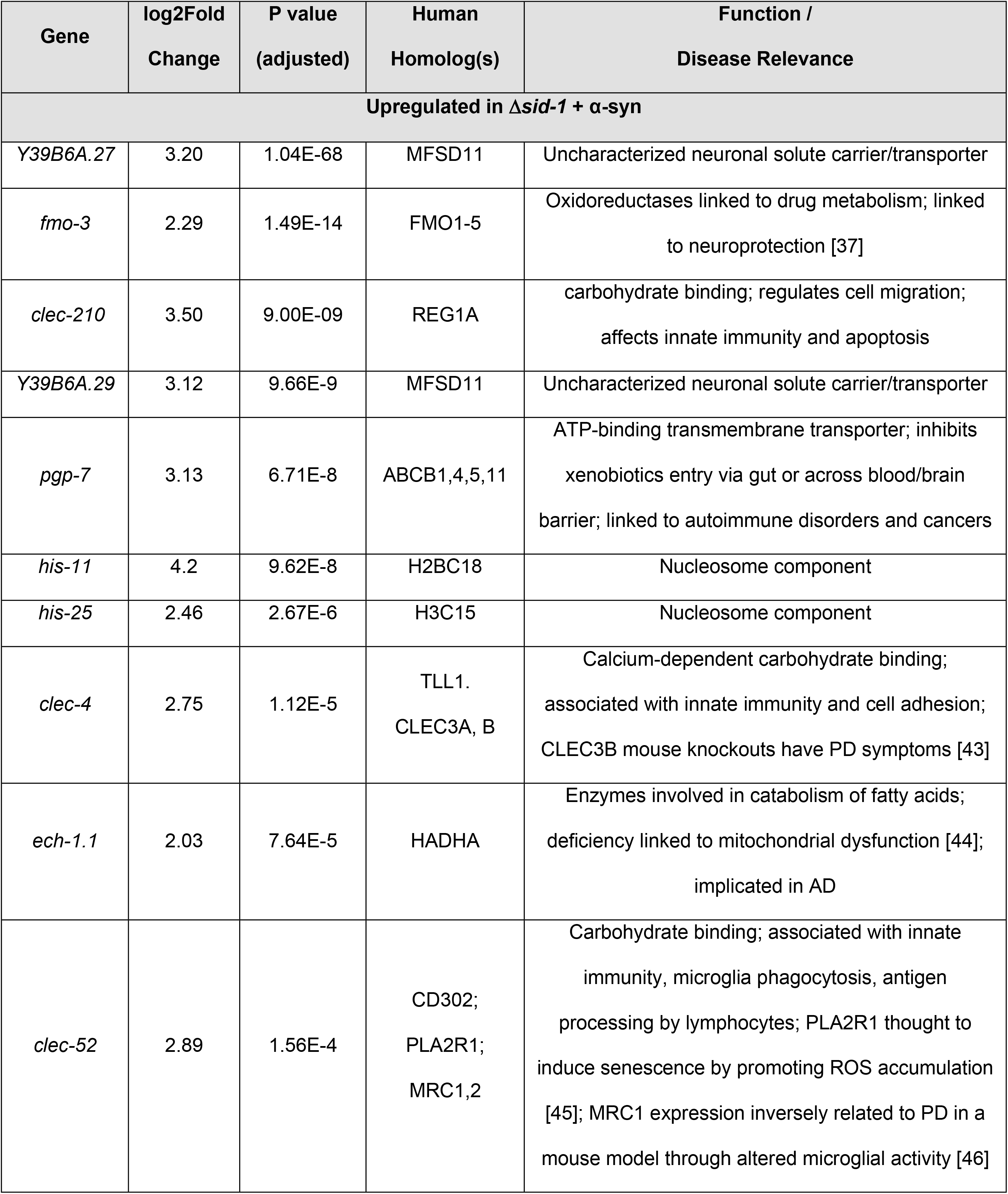
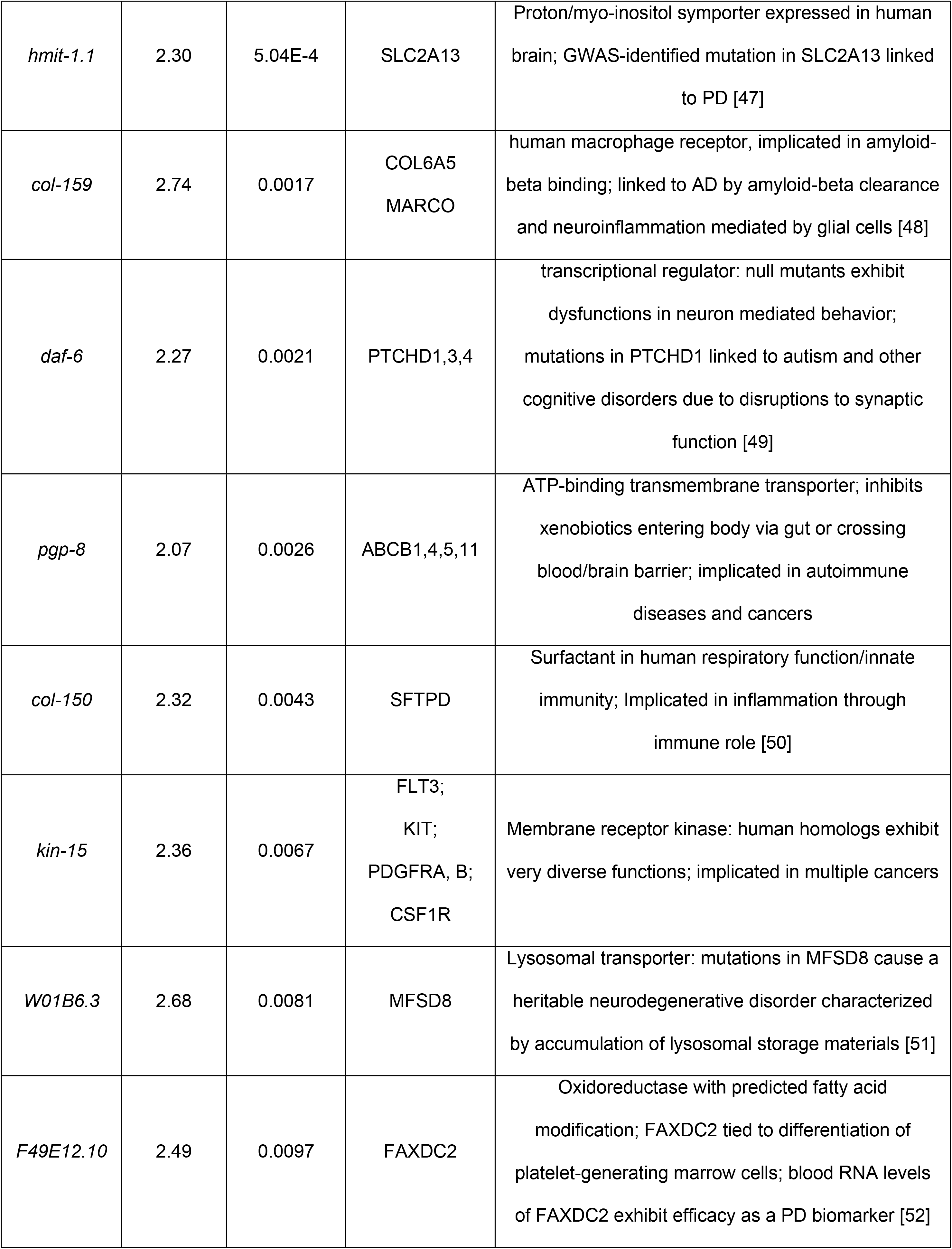
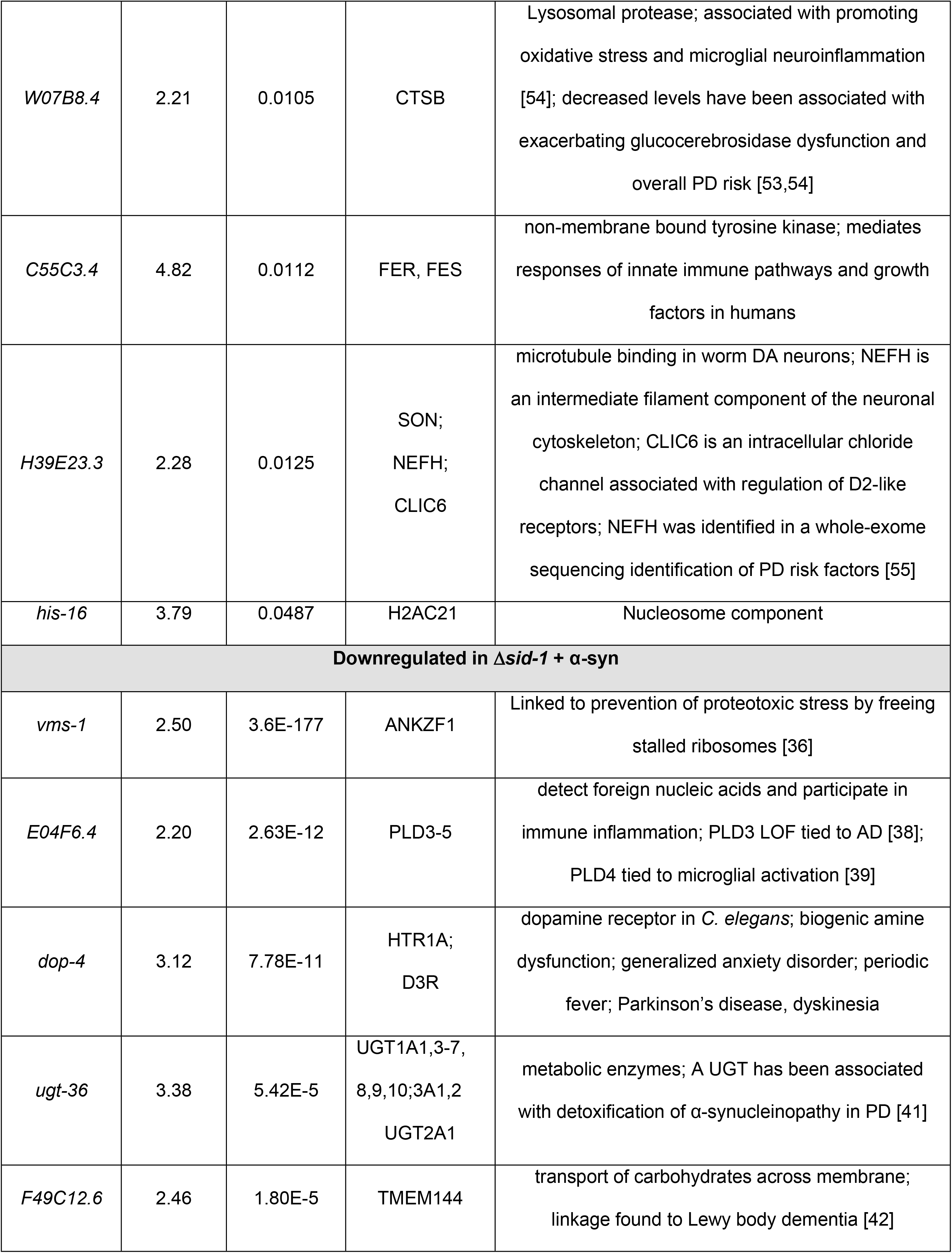
Differentially Expressed Genes (DEGs): α-syn vs. α-syn + Δ*sid-1*

### Combined loss of both SID-1 and PGP-8 function reduces *mir-2* overexpression-induced enhancement of neurodegeneration

The transcriptomic comparison of α-syn worms vs. α-syn worms in the *sid-1* mutant background revealed two DEGs that code for transmembrane transporter proteins that were upregulated in the *sid-1* mutants compared to the α-syn background alone: *pgp-7* and *pgp-8*. These are P-glycoprotein (P-gp) related genes with have human homologs that are ATP-binding cassette (ABC) transporters. This class of transmembrane transporters have mostly been found to function as exporters and are involved in the efflux of toxins, lipopeptides and lipophilic drugs in eukaryotes, including humans [56,57,58]. However, exceptions have been reported, as the polarity of P-gp transport is neither exclusively unidirectional nor is it limited to efflux.

Animals mutant for *sid-1* in the α-syn background exhibited significantly higher levels of *pgp-8* mRNA transcripts (>2-fold; p =0.0026) compared to the α-syn background alone (with wild-type *sid-1*) (Fig. 5C). Given the availability of a *pgp-8* deletion mutant, we reasoned it worthwhile to investigate the prospect that the upregulation of *pgp-8* was part of a potential compensatory response to systemic SID-1 deficiency. Therefore, α-syn*^mir-2^* ^OE gut^ worms were crossed to animals mutant for *pgp-8*, as well as to *pgp-8*; *sid-1* double mutants and were subsequently scored for any changes in α-syn-induced DA neurodegeneration. When *pgp-8* alone was mutant, intestinal *mir-2* overexpression significantly enhanced neurodegeneration compared to α-syn only worms (Fig. 5D). However, when *pgp-8* and *sid-1* were both mutant, the degenerative effect of intestinal *mir-2* overexpression was negated (Fig. 5D). In the absence of *mir-2* overexpression, neither *pgp-8* mutants or *pgp-8*; *sid-1* double mutants exhibited any significant alterations in α-syn-induced DA neurodegeneration (Fig. 5E, Fig. 5F). These results suggested that the absence of both SID-1 and PGP-8 function disrupt cellular entry of *mir-2* and consequentially impedes the silencing of the neuroprotective targets of *mir-2*, as opposed to what was observed when either *sid-1* or *pgp-1* were independently mutant (Fig. 5L).

Having identified a requirement for PGP-8 function in the ability of *mir-2* to modulate DA neurodegeneration in the absence of functional SID-1, we wanted to explore the prospect that PGP-8 may be functioning as an importer, as opposed to the more typical role for P-gps in efflux of a broad range of substrates from within cells. To examine this, we exposed N2 wild-type worms and worms with the *pgp-8* mutation to rhodamine 123, a fluorescent dye that localizes to mitochondria and is a known substrate for P-gp ABC transporters [59]. The quantified average pixel intensity of rhodamine 123-induced fluorescence displayed by *pgp-8* mutant worms was noticeably and significantly decreased compared to that of N2 worms exposed to this dye (Fig. 5G-I). These data are indicative of a previously uncharacterized role for PGP-8 in the influx of substrates. To explore this further, we conducted a similar set of experiments using AIM-100 to inhibit SID-3 in worms in the α-syn*^mir-2^* ^OE gut^ background that also had either the *pgp-8* mutation alone or both *pgp-8* and *sid-1* mutated. When only *pgp-8* was mutant (solvent group), *mir-2*-induced enhancement of α-syn-mediated DA neurodegeneration was observed (Fig. 5J). In contrast, when AIM-100 was administered to these same worms, α-syn-mediated neurodegeneration was attenuated (Fig. 5J). This implies that loss of PGP-8 function is of little consequence in the presence of functional SID-1, as AIM-100 treatment exerts the same neuroprotective effect on *mir-2*-induced neurodegeneration as was previously observed (Fig. 4B, Fig. 4C). When both *pgp-8* and *sid-1* were mutant in the α-syn*^mir-2^* ^OE gut^ background (solvent group), the DA neurodegeneration observed was consistent with the effect seen with untreated α-syn animals containing both *pgp-8* and *sid-1* mutations: there was no significant alteration in neurodegeneration (Fig. 5K). When AIM-100 was administered to *pgp-8*; *sid-1* double mutants, intestinal *mir-2* overexpression had no additive effect on α-syn-induced DA neurodegeneration (Fig. 5K). This indicates that the inhibition of SID-3 activity caused by AIM-100 attenuates the cell non-autonomous neurodegenerative effect of *mir-2* overexpression only when *sid-1* is intact (wild-type), and this inhibition is independent of *pgp-8* status. Taken together, the transcriptional response to *sid-1* depletion clearly exerts a functional change in how dsRNAs like *mir-2* (and likely other miRNAs) impart their effects on gene silencing (Fig. 5L). In conjunction with the observed dependence on *pgp-8* for intestinal *mir-2* overexpression to cause neurodegeneration, these results collectively suggest that an alternative pathway, independent of *sid-1*, may be plausibly involved in the epigenetic modulation of DA neurodegeneration.

## Discussion

Whereas precise causes of PD are complex and unresolved, this movement disorder is unequivocally a combined consequence of genetic predisposition and epigenetic response to exogenous (i.e., toxins) and/or intrinsic environmental (i.e., microbiome) factors. It is to be expected that an increased susceptibility to neurodegenerative states accompanying a chronic source of cellular stress, such as expression of a chromosomally integrated multicopy α-syn transgene, further exacerbated with time and aging, evokes inherent organismal responses. Indeed, numerous studies across a wide range of cellular and animal systems have examined the transcriptional response to overexpression of α-syn or other genes, mutations in those loci, various PD-associated toxins, addition of pre-formed α-syn fibrils, and more. None of these models enable rigorous experimentation aimed at understanding the consequences on neurodegeneration that may arise from disruption and changes in the efficacy of small RNA transport and the systemic effects of gene silencing, as has proven possible using *C. elegans* [21]. Recent results of transcriptional profiling experiments using a different *C. elegans* α-syn model documented extensive changes in the expression of miRNAs and piRNAs [16]. Our present study diverges from further descriptive cataloging of α-syn-induced gene expression to instead consider a different, more mechanistic question. Specifically, how is it that small RNAs exert a functional impact as modifiers of the degeneration of DA neurons?

Here we have demonstrated that the DA neurons of *C. elegans* are receptive and responsive to epigenetic regulation by endogenous, exogenous, cell autonomous, and cell non-autonomous sources of dsRNA. In discerning that established components of the systemic gene silencing machinery of *C. elegans*, SID-1 and SID-3, can function as effectors of α-syn-induced DA neurodegeneration, we provide a roadmap for future investigation of conserved targets of their activity in modulating neurodegenerative disease, including numerous previously unrecognized neuroprotective gene products.

It is important to note that even in *sid-1* mutants, lacking systemic dsRNA transporter function, the process of endogenous gene silencing appears to remain intact within cells that themselves transcribe dsRNAs. Thus, the abolishment or inhibition of gene silencing in *sid-1* or *sid-3* mutants only applies to genes that are silenced by mobile dsRNAs that originate outside of target cells and require transport into the cytoplasm. Moreover, it appears as though the neuroprotective benefit DA neurons receive from reduced gene silencing in α-syn worms is largely dependent on SID-1 transporter function. Contrary to the prevailing dogma in the *C. elegans* field [60], the levels of SID-1 present in the DA neurons are clearly not too low to exert a functional effect that can be revealed either by *sid-1* knockout and/or dsRNA expression from within the animal. While neuronal *sid-1* expression levels are unequivocally lower on average when compared to other cell types, recent data that have emerged from the *C. elegans* Neuronal Gene Expression Map & Network (CeNGEN) consortium showing that *sid-1* is expressed in DA neurons [61]. Perhaps the comparatively limited levels of basal *sid-1* expression are reflected by the observation that, in *sid-1* mutant animals, the only significant reduction in neurodegeneration was more limited to later in life, when the progressive effect of α-syn-induced DA neurodegeneration is typically most extensive. However, *sid-1* mutants still exhibit significantly improved behavioral functionality at earlier timepoints, and as demonstrated by the restoration of DA neurotransmission via the BSR.

The protection from α-syn neurotoxicity observed in *sid-3* mutants was more robust at all ages examined. SID-1 is still present in these animals, but it is subjected to increased endocytic recycling at the plasma membrane in the absence of SID-3 non-receptor tyrosine kinase activity. The difference in neuroprotection observed in *sid-3* mutants in comparison to *sid-1*; *sid-3* double mutants, specifically at timepoints when *sid-1* mutants alone showed little or no neuroprotection, is not unexpected. This is likely a reflection of SID-3 also having an important SID-1-and dsRNA-independent function as a regulator of the basal endocytosis of the *C. elegans* dopamine transporter, DAT-1. As was previously described, ACK1 (the human ortholog of SID-3) regulates internalization of DAT on the plasma membrane of DA neurons to modulate synaptic dopamine levels and neurotransmission [25]. Accordingly, others and we have also reported that *dat-1* mutants in *C. elegans* protect DA neurons from degeneration [62, 63]. Whereas *dat-1* gene expression is limited to the DA neurons, the *sid-3* gene is more broadly expressed, as is *sid-1.* Thus, it is tempting to speculate that this duality of transporter regulation observed within the DA neurons is a unique evolutionary confluence of function inherent to SID-3. Yet, it is equally interesting to consider other partners of SID-3 endocytic regulation, especially given the range of substrates ACK1 is known to phosphorylate in mammalian cells, where it is implicated in tumor survival in a variety of cancers [64]. DA biosynthesis, packaging, synaptic release and reuptake, as well as its reception, metabolism, and function are collectively well established as a tightly regulated integration of evolutionarily conserved processes. Therefore, the endocytic control of both neurotransmitter and dsRNA transporter function by SID-3 represents a nexus of genetic and epigenetic convergence that promotes a capacity for dynamic responsivity to both internal and external influences on an organism.

The comparative transcriptomic analysis performed using transgenic α-syn animals either with or without the *sid-1* mutation provides a window into putative candidate gene products that exert a differential function when SID-1 dsRNA transporter activity is absent. In this regard, among the list of 27 DEGs with human orthologs identified, the increased expression of the *pgp-7* and *pgp-8* genes were of particular interest. Both these genes encode highly conserved members of the P-glycoprotein (P-gp) ABC transporter family, primarily known for their broad substrate specificity in the translocation of a variety of molecules across the plasma membrane of cells. P-gp transporters are generally classified as facilitating the efflux of toxins, drugs, lipophilic peptides, and other small molecules out of cells. Nevertheless, examples of ABC transporters functioning to import substrates exist and are present in both plants and animals, including humans [65, 66]. Indeed, even the archetypical P-gp, the MDR1/ABCB1 multidrug resistance transporter, has been shown to function in the influx of substrates from the epidermis to the skin of mice, whereas *mdr1 (-/-)* mice lack the deposition of epidermally derived molecules [59]. In a feat of protein engineering, it was recently demonstrated that the directionality of transport could be reversed from efflux to influx by mutagenesis of conserved residues in MDR1/ABCB1 [67]. Importantly, among the earlier mechanistic observations on RNAi in *C. elegans* was a report that demonstrated that the efficacy of dsRNA gene silencing showed a requirement for members of the P-gp ABC transporter family to be expressed in the same cells as the gene being targeted for knockdown [68]. Through a screen of 43 different ABC transporters for which mutants had been isolated, it was determined that a total of 9 (about 30%) of these genes non-redundantly influenced RNAi activity. The *C. elegans* genome encodes over 60 members of the ABC transporter family, most of which have not been extensively characterized with respect to function, including PGP-8, which was not among the genes tested at the time of this prior study, but it has been reported to exhibit an enriched expression in neurons [69]. Interestingly, over 100 genes have been shown to play a role in the network of genes comprising RNAi function in *C. elegans,* yet the SID proteins and these ABC transporters are among very few of those that localize to membranes [70]. To the best of our knowledge, no precedent exists for the translocation of small RNAs by ABC transporter proteins, and we are reluctant to speculate that our observations are a result of such a mechanism. Whatever substrate(s) of PGP-8 are involved with its effect on either the systemic spread of RNAi or in a more localized role in DA neurons, this relationship merits further attention.

Still, if specific gene products functionally compensate for SID-1 in its absence, then dsRNA import into cells would be lost, or at least diminished when they too were mutant. It was therefore notable that the combined genetic depletion of both *sid-1* <u>and</u> *pgp-8* in double-mutant animals abolished the ability of an intestinally expressed miRNA, *mir-2,* to significantly enhance neurodegeneration. Whereas when either *sid-1* or *pgp-8* were individually mutant, the cell non-autonomous overexpression of *mir-2* had no influence on α-syn-induced DA neurodegeneration. Arguably, the most fascinating result from this study was that, even in a *sid-1* mutant background, intestinal expression of *mir-2* was still able to enhance the degeneration caused by α-syn in DA neurons. Taken on its own, this could suggest that *mir-2* is silencing target genes locally, in the intestinal cells, which would then indirectly affect DA neuron degeneration. This would be an appealing explanation, as there is a growing precedent for gut-to-neuron signaling as being a contributing factor in the etiology of PD [71, 72]. In our case, this possibility is contested since AIM-100 administration, which decreases dsRNA import in all cells expressing *sid-3*, abolishes the ability of *mir-2* to enhance α-syn-mediated neurodegeneration. Furthermore, as it is known that SID-1 can import dsRNAs of at least up to 500 basepairs, overexpression of *mir-2* (pre-miRNA, 98bp) should be readily transported by SID-1 [73]. Taking this into account, this result remains enigmatic, since the ability of *mir-2* to access cells, silence target genes and subsequently affect α-syn-mediated neurodegeneration should be completely dependent on the presence of functional SID-1. Considering that knockout of <u>both</u> *pgp-8* and *sid-1* (*pgp-8*; *sid-1* double mutants) eliminates the enhanced neurodegeneration caused by intestinal *mir-2* overexpression, this may provide a clue to the organismal dynamics that are at play when SID-1 transport activity is compromised or depleted by mutation.

Since the transmission of *mir-2* appears to induce an increased susceptibility to neurodegeneration, this counterintuitively suggests that the basal expression levels of certain genes (i.e., targets of miRNAs) are maintained in a state that hinders the capacity of DA neurons to protect themselves. However, that premise represents an oversimplification of the integrated dynamics of the various cellular subcomponents involved in maintaining neuronal homeostasis. First, the enzymatic pathways involved in the biosynthesis of DA, as well as the cellular machinery responsible for vesicular packaging, trafficking, metabolism, synaptic release and reuptake are all tightly regulated processes conserved between worms and humans. Second, while the misfolding and oligomerization of α-syn, along with its concomitant impact on cellular dynamics are central to the underlying pathology of PD [74], the fact that the *C. elegans* genome does not naturally contain an α-syn homolog complicates the interpretation of why a loss or decrease of dsRNA-mediated gene silencing is beneficial in this context. Third, the levels of DA itself play a vital role in the selective vulnerability of DA neurons to a variety of potential sources of damage (i.e., oxidative, mitochondrial, lysosomal). In the case of PD, the critical dosage-dependent misfolding and oligomerization associated with α-syn neurotoxicity introduces a significant impediment to neuronal homeostasis. This imbalance is further compounded by *in vivo* evidence from both *C. elegans* and mouse PD models that demonstrated a direct role for the dysregulation of DA levels in neurodegeneration because of the interaction of this neurotransmitter with α-syn [75]. Therefore, optimal dopaminergic neuronal functionality is dependent on a suite of factors that, if perturbed by any of a series of environmental, genetic, or epigenetic factors, can profoundly impact susceptibility to neurodegeneration. In considering the pivotal roles that both SID-1 and SID-3 have in the orchestration of epigenetic regulation by small RNAs, systemically in *C. elegans*, it is not difficult to imagine the impact on DA neuron homeostasis and survival that we have observed.

This study offers a new perspective that positions the activity of SID-1 and SID-3, and their implicit regulation of epigenetic gene silencing, in a pivotal relationship with neurodegeneration, specifically as it pertains to synucleinopathies like PD. While it is likely to vary substantially, the conceptual model that continues to emerge in this research may prove applicable to other cells and neuron subtypes in *C. elegans*. Furthermore, although stark anatomical differences and other complexities obviously preclude direct comparisons between organismal mechanisms of small RNA transport among highly diverse species, the existence of human orthologs of both SID-1 and SID-3 points to mechanistic commonalities that are yet to be discerned [30,31,76]. Likewise, even though the conservation of miRNA sequences and structures is limited between humans and worms, this does not nullify the value in being able to identify evolutionarily conserved genes that are targets of epigenetic regulation, and whose expression is influenced by the dsRNA transport machinery. Overall, this approach represents a strategy to reveal functional modifiers and therapeutic targets previously unrecognized for their neuroprotective capacities. Further investigation toward fine tuning their expression as individual targets or, perhaps, as co-regulated factors with a combined potency, should be explored with the requisite sense of urgency needed to implement innovations with potential therapeutic benefit for neurodegenerative diseases.

## Materials and Methods

### C. elegans strains

Experimental nematodes were reared and maintained on OP50-1 *E. coli* bacteria (unless otherwise noted) at 20°C under standard laboratory conditions [77]. The following strains were obtained from the *Caenorhabditis* Genetics Center: N2, CB1112 (*cat-2*(e1112)), RB1916 (*pgp-8(ok2489)*) and HC46 [ccls4251(myo-3::GFP-NLS (nuclear localized), myo-3::GFP-MITO (mitochondrial localization); mls11(myo-2::GFP)]. Integrated transgenic strain BY250 (*vtIs7* [P*_dat-1_::*GFP]) was a gift from Randy Blakely (Florida Atlantic Univ.). Integrated transgenic lines crossed into BY250 include: UA423 (vtIs7[P*_dat-1_*::GFP]; *sid-1*(pk3321). There were 3 separate integrated transgenic α-syn neurodegeneration models that were used in this study: UA44 (*baIn11*[P*_dat-1_*:: α-syn (human, wild-type), P*_dat-1_*::GFP]), UA372 (*baIn54*[P*_dat-1_*::α-syn (human, A53T mutation), P*_unc-54_*::tdTomato]), and UA196 (*baIn11*[P*_dat-1_*:: α-syn (human, wild-type), P*_dat-1_*::GFP]; *baIn33* [P*_dat-1_*::*sid-1*, P*_myo-2_*::mCherry]; *sid-1*(pk3321)).

Integrated lines crossed into UA44 include: UA415 (*baIn11*[P*_dat-1_*::α-syn (human, wild-type), P*_dat-1_*::GFP]; *sid-1*(pk3321)), UA416 (*baIn11*[P*_dat-1_*::α-syn (human, wild-type), P*_dat-1_*::GFP]; *sid-3*(ok973)), UA414 (*baIn11*[P*_dat-1_*::α-syn (human, wild-type), P*_dat-1_*::GFP]; *sid-1*(pk3321); *sid-3*(ok973)), UA418 (*baIn11*[P*_dat-1_*::α-syn (human, wild-type), P*_dat-1_*::GFP]; *mir-2*(gk259)), UA417 (*baIn11*[P*_dat-1_*::α-syn (human, wild-type), P*_dat-1_*::GFP]; *sid-1*(pk3321); *mir-2*(gk259)), UA419 (*baIn11*[P*_dat-1_*::α-syn (human, wild-type), P*_dat-1_*::GFP]; *pgp-8*(2489)), and UA436 (*baIn11*[P*_dat-1_*::α-syn (human, wild-type), P*_dat-1_*::GFP]; *sid-1*(pk3321); *pgp-8*(2489). Integrated transgenic lines crossed into UA372 include: UA420 (*baIn54*[P*_dat-1_*:: α-syn (human, A53T), P*_unc-54_*::tdTomato]; *sid-1*(pk3321)), and UA421 (*baIn54*[P*_dat-1_*:: α-syn (human, A53T), P*_unc-54_*::tdTomato]; *sid-3*(ok973)). Integrated transgenic lines crossed into UA196 include: UA437 (baIn11[[P*_dat-1_*:: α-syn (human, wild-type), P*_dat-1_*::GFP]; baIn33[P*_dat-1_*::*sid-1*, P*_myo-2_*::mCherry]; *sid-1*(pk3321); *mir-2*(gk259)). Stable transgenic lines crossed into UA44 include: UA273 (*baIn11*[P*_dat-1_*::α-syn (human, wild-type), P*_dat-1_*::GFP]; *baEx161*[P*_dat-1_*::*mir-2* (pre-miRNA), P*_unc-54_*::mCherry)]), UA408 (*baIn11*[P*_dat-1_*::α-syn (human, wild-type), P*_dat-1_*::GFP]; *baEx226*[P*_ges-1_*::*mir-2* (pre-miRNA), P*_unc-54_*::tdTomato]), UA409 (*baIn11*[P*_dat-1_*::α-syn (human, wild-type), P*_dat-1_*::GFP]; *baEx226*[P*_ges-1_*::*mir-2* (pre-miRNA), P*_unc-54_*::tdTomato]; *mir-2*(gk259)), UA410 (*baIn11*[P*_dat-1_*::α-syn (human, wild-type), P*_dat-1_*::GFP]; *baEx226*[P*_ges-1_*::*mir-2* (pre-miRNA), P*_unc-54_*::tdTomato]; *sid-1*(pk3321)), UA411 (*baIn11*[P*_dat-1_*::α-syn (human, wild-type), P*_dat-1_*::GFP]; *baEx226*[P*_ges-1_*::*mir-2* (pre-miRNA), P*_unc-54_*::tdTomato]; *mir-2*(gk259); *sid-1*(pk3321)), UA412 (*baIn11*[P*_dat-1_*::α-syn (human, wild-type), P*_dat-1_*::GFP]; *baEx226*[P*_ges-1_*::*mir-2* (pre-miRNA), P*_unc-54_*::tdTomato]; *pgp-8*(ok2489)), and UA413 (*baIn11*[P*_dat-1_*::α-syn (human, wild-type), P*_dat-1_*::GFP]; *baEx226*[P*_ges-1_*::*mir-2* (pre-miRNA), P*_unc-54_*::tdTomato]; *pgp-8*(ok2489); *sid-1*(pk3321)).

### Transgenic Line Construction

All stable transgenic lines were created by injecting both the transgene and co-injection marker at a concentration of 50 ng/μL. All stable transgenic lines were created by injecting into UA44 worms and then subsequently crossing into indicated mutant backgrounds, except UA272 which was first injected into N2 worms and then subsequently crossed into the UA44 background. A Gateway Entry Clone consisting of the sequence corresponding to the precursor miRNA (pre-miRNA) for *mir-2* with sequence: 5’ - TAAACAGTATACAGAAAGCCATCAAAGCGGTGGTTGATGTGTTGCAAATTATGACTTTCATATCACAGCCAGCTTTGATGTGCTGCCTGTTGCACTGT – 3’ was generated using these primers (includes Gateway Tail Sequences):

Forward: 5’ - GGGGACAAGTTTGTACAAAAAAGCAGGCTCCTAAACAGTATACAGAAAGCCATCAA AGC

Reverse: 5’ – GGGGACCACTTTGTACAAGAAAGCTGGGTCACAGTGCAACAGGCAGCACATC – 3’.

Gateway Expression Clones including either the *dat-1* or *ges-1* promoter sequence directly upstream of the pre-*mir-2* sequence were generated from this Gateway Entry Clone. Gateway Expression Clones consisting of the *unc-54* promoter sequence directly upstream of the sequences corresponding to either the fluorescent protein tdTomato or mCherry were used as co-injection markers to generate stable transgenic lines.

### Dopaminergic Neurodegeneration Analysis

Worms were age synchronized by performing 3-5-hour egg lays. All worms were kept at 20°C for the duration of each experiment. When analyzing stable transgenic lines, great care was taken to isolate and test only worms with the transgenic marker. The extent of neurodegeneration was determined for each replicate by placing worms in a 6 μL drop of 10 mM levamisole (dissolved in 0.5x S Basal buffer) on a glass coverslip. This drop of levamisole on the coverslip was inverted and placed on a 2% agarose pad made on a microscope slide, immobilizing worms to aid in visualization. Using a Nikon Eclipse E600 epifluorescence microscope with the addition of a Nikon Intensilight C-HGFI fluorescent light source, the 6 DA neurons in the head region of worms (4 CEPs and 2 ADEs) were scored for the extent of neurodegeneration by using GFP fluorescence as a proxy. Each worm was scored as either normal or degenerative. Worms were considered normal only if all 6 DA neurons in the head region had completely intact cell bodies and dendritic processes. Worms were considered degenerative if at least 1 out of the 6 DA neurons was absent, had broken dendritic processes, or exhibited cell body abnormalities. Experiments using integrated transgenic lines consisted of 3 biological replicates, and each biological replicate consisted of 30 worms. Experiments using stable transgenic lines without any drug administration consisted of 3 biological replicates (3 separate stable lines), apart from strain UA272 which consisted of 2 biological replicates. Each biological replicate consisted of 1 stable line of each strain, each consisting of 3 technical replicates. Each technical replicate consisted of 30 worms, resulting in a total of 90 worms per biological replicate. AIM-100 experiments using stable transgenic lines consisted of 3 biological replicates (3 separate stable lines). Each biological replicate consisted of 1 stable line of each strain, each consisting of 1 technical replicate. Each technical replicate consisted of 30 worms, resulting in a total of 30 worms per biological replicate. Statistical significance between groups was determined with GraphPad Prism Software.

### Neuron Image Acquisition

Images of DA neurons were obtained by placing worms in a 6 μL drop of 10 mM levamisole (dissolved in 0.5x S basal Buffer) on a glass coverslip. This drop of levamisole on the coverslip was inverted and placed on a 2% agarose pad made on a microscope slide, immobilizing worms to aid in visualization. Fluorescence microscopy was performed with a Nikon Eclipse E800 epifluorescence microscope equipped with an Endow GFP HYQ filter cube. Images were captured using a Cool Snap CCD camera (Photometrics) with Metamorph software (Molecular Devices).

### Basal Slowing Response (BSR) Assays

BSR assays were conducted similarly as reported by Martinez and colleagues [78]. The BSR assay was performed on day 4 post-hatching and worms were reared on OP50-1 *E. coli*. N2 and CB1112 (*cat-2*(e1112)) strains were used to validate the assay: N2 worms exhibit a normal BSR and *cat-2* mutants exhibit a defective BSR. Ring plates (NGM) with a 4 cm outer ring of HB101 *E. coli* (OD_600_ of 0.6-0.7 A) and a 1 cm inner circle (unseeded) were used. Individual worms were washed of native bacteria by allowing them to thrash briefly in a 10 μL drop of 0.5 x S basal buffer. Single worms were placed in the center of ring plates in the unseeded portion, upon which time video recording was initiated. Locomotion was recorded using an automated video tracking system (MBF Bioscience) and analyzed using WormLab Software (Version 4.0.5; MBF Bioscience). Videos were taken of individual worms, each on a separate plate. The average peristaltic speed (μm/second) was recorded of the 100 frames before the head of the worm touched the bacterial boundary on the unseeded portion of the ring plate, and the 100 frames after the head touched the bacterial boundary. Three replicates were performed with each strain, each replicate consisting of ten individual worms. The average peristaltic speed on food was compared to the average peristaltic speed off food and converted to a ratio (on/off) that was inverted and normalized to the N2 value to represent BSR as a percentage response compared with N2, which was defined as 100%. Statistical significance between groups was determined with Graphpad Prism Software.

### RNAi Experiments: Plates and Bacterial Growth Conditions

RNAi plates were made by adding ampicillin and Isopropyl β-D-1 thiogalactopyranoside (IPTG) at a final concentration of 100 μg/mL and 1 mM respectively to NGM. RNAi bacteria (HT115 *E. coli*) containing either the empty L4440 feeding vector or this same vector containing sequences anti- to target genes desired to be knocked down were grown in LB media with the addition of ampicillin at a final concentration of 100 μg/mL at 37°C while shacking for 16 hours. RNAi bacteria was seeded onto RNAi plates and allowed to fully dry and adhere to the plates in a biological safety cabinet. The resulting plates were then incubated at 20°C overnight to allow for dsRNA induction.

### AIM-100 Drug Administration

AIM-100 plates were made by first dissolving AIM-100 (Tocris Cat. No. 4946) in a molecular grade 200 proof ethanol stock and then adding to NGM at a final concentration of 100 μM AIM-100 / 0.1% ethanol. Control (solvent) plates contained the same concentration of ethanol (0.1%) as AIM-100 plates, but without the addition of AIM-100. All AIM-100 plates used for experiments were seeded with OP50 *E. coli*.

### Rhodamine 123 Administration and Pixel Intensity Analysis

Rhodamine 123 plates were made by first dissolving rhodamine 123 (VWR Cat. No. 89139-378) in dimethyl sulfoxide (DMSO) and then adding to NGM at a final concentration of 10 μM. Plates were seeded with OP50 *E. coli* after solidification and used for rhodamine 123 treatment immediately. Images of whole worms 48 hours post-hatching were obtained by placing worms in a 6μL drop of 10 mM levamisole (dissolved in 0.5x S basal Buffer) on a glass coverslip. This drop of levamisole on the coverslip was inverted and placed on a 2% agarose pad made on a microscope slide, immobilizing worms to aid in visualization. Fluorescence microscopy was performed with a Nikon Eclipse E800 epifluorescence microscope equipped with an Endow GFP HYQ filter cube. Images were captured using a Cool Snap CCD camera (Photometrics) with Metamorph software (Molecular Devices). 30 images per replicate were captured per genotype, and each genotype consisted of 3 independent replicates. Once imaged, worms were analyzed for average pixel intensity of rhodamine 123-induced fluorescence using a standardized 50-pixel-diameter circle at the midpoint of the worm bodies using the vulva as an anatomical marker, using the Metamorph software. Representative images of each genotype were taken in the same fashion.

### Transcriptomic Analysis

RNA was isolated from strains UA44 and UA415 at day 6 post-hatching. 3 separate replicates were isolated for each strain. Large quantities of worms were obtained by performing 2-hour egg lays with 20-30 gravid adults on large NGM plates seeded with OP50-1 *E. coli*. All worms destined for RNA isolation were grown at 20°C. Starting at day 4 post-hatching, 400 worms were transferred to 100 mm fresh seeded plates to prevent starvation and mixing of target worms and progeny. At day 6 post-hatching, for each strain, for each replicate, 380 worms were transferred to an unseeded NGM plate. Great care was taken to only transfer target worms and to avoid carryover of embryos. Worms were washed off unseeded plates with 0.5x M9 into sterilized glass conical tubes. Worms were washed 4 times with 0.5x M9 to reduce bacteria in samples. After the 4^th^ wash, worms were purged for 20 minutes to reduce the number of bacteria in the gut of the worms, and this was done by rocking the tubes back and forth on a nutator. Worms were then washed 2 times with double-distilled deionized water to rid of any remaining bacteria. Worms were then transferred to 1.5 mL RNAse-free lo-bind microcentrifuge tubes and as much supernatant was removed as possible. 500 μL of Trizol (ThermoFisher Scientific) was then added to each tube, and then immediately vortexed for 30 seconds and placed in liquid nitrogen until frozen completely. Tubes were then thawed at 37°C. This freeze/thaw process was then performed the same way 6 more times. After the last thaw, all tubes were vortexed for 30 seconds and then put on ice for 30 seconds. This vortex/ice process was then performed the same way 7 more times. All tubes were then allowed to incubate at room temperature for 5 minutes. Then 100 μL of chloroform was added to each tube, and each tube was immediately inverted continuously for 15 seconds. The tubes were then allowed to incubate at room temperature for 5 minutes to allow for phase separation. The tubes were then centrifuged for 15 minutes at 4°C at 12,000 rpm. The top aqueous phase from each tube was then transferred to new 1.5 mL RNAse-free microcentrifuge tubes, and an equal volume of RNAse-free 70% ethanol was added to each tube and mixed by gentle inversion. At this point, the aqueous phase mixed with ethanol was subjected to the protocol outlined in the Qiagen RNeasy Micro Kit, and reagents and materials from this kit were used for the remainder of the isolation. Ultimately, RNA was eluted with 35 μL of RNAse-free water.

RNA samples were sent to and sequenced by the Novogene Corporation, Inc.; Sacramento, CA. Sequencing was eukaryotic mRNA sequencing. Novogene also performed bioinformatic analysis to determine DEGs, along with transcript levels, log2fold changes, and adjusted and non-adjusted p-values. An Illumina Novaseq platform was used for a paired-end 150 basepair sequencing strategy (short reads) to sequence cDNA libraries corresponding to RNA samples. Transcript expression levels were determined using the RPKM (Reads Per Kilobases per Million reads) method [79]. Data was filtered by removing adaptors and low-quality reads. Bioinformatic analysis was done using HISAT2, version 2.1.0-beta [80], with a parameter of “mismatch=2” to map samples to the N2 reference genome (ftp://ftp.ebi.ac.uk/pub/databases/wormbase/parasite/releases/WBPS13/species/caenor habditis_elegans/PRJNA13758/caenorhabditis_elegans.PRJNA13758.WBPS13.genomi c.fa.gz). The resultant BAM files from HISAT2 were input for quantification of reads with HTSeq, version 0.6.1, with parameter “-m union” [81]. Differentially expressed genes (DEGs) were extracted using DESeq2 version 1.10.1 in R [82]. Significance was determined if the p-adjusted value was less than 0.057. DESeq2 utilized a Benjamini-Hochberg method of normalization to calculate the adjusted p-value, a powerful method for controlling the false discovery rate. Heatmaps were created in R using package “pheatmap” (Kolde, 2015). Gene ontology (GO) analyses were created using WormCat bioinformatics resource [84]. Genes that were significantly up- or down-regulated as determined by an adjusted p-value of less 0.05 were used in the analysis.

### Statistical Analysis

All statistical analysis was performed with GraphPad Prism Software (Version 9.0.0). When analyses were performed with data containing multiple days, comparing every group to every other group, a Two-Way ANOVA was used with Tukey’s *post hoc* analysis. When analyses were performed with data containing multiple days, comparing only groups within the same day to each other, a Two-Way ANOVA was used with a Šídák’s *post hoc* analysis. When analyses were performed with data containing one single day, and there were only two groups, an unpaired, two-tailed t-test was used. When analyses were performed with data containing one single day, with more than two groups and only comparing back to a control, a One-Way ANOVA was used with Dunnett’s *post hoc* analysis. All data assessed involving AIM-100 administration used a Two-Way ANOVA with Tukey’s *post hoc* analysis.

## Acknowledgements

We would like to thank the past and present members of the Caldwell lab for their collegiality and helpful discussions, including Brian Smithers, Paige Dexter, and Susan DeLeon for their technical assistance. Special thanks to Dr. Janna Fierst for her time and expert assistance with bioinformatic analyses. This paper is dedicated to the memory of our brilliant and beloved student, colleague and friend, Dr. Susan M. DeLeon-Gulbronson.

